# Robust and Accurate Bayesian Inference of Genome-Wide Genealogies for Large Samples

**DOI:** 10.1101/2024.03.16.585351

**Authors:** Yun Deng, Rasmus Nielsen, Yun S. Song

**Affiliations:** Center for Computational Biology, University of California, Berkeley, USA; Department of Statistics, University of California, Berkeley, USA; Department of Integrative Biology, University of California, Berkeley, USA; Center for GeoGenetics, University of Copenhagen, Denmark; Computer Science Division, University of California, Berkeley, USA

## Abstract

The Ancestral Recombination Graph (ARG), which describes the full genealogical history of a sample of genomes, is a vital tool in population genomics and biomedical research. Recent advancements have increased ARG reconstruction scalability to tens or hundreds of thousands of genomes, but these methods rely on heuristics, which can reduce accuracy, particularly in the presence of model misspecification. Moreover, they reconstruct only a single ARG topology and cannot quantify the considerable uncertainty associated with ARG inferences. To address these challenges, we here introduce SINGER, a novel method that accelerates ARG sampling from the posterior distribution by two orders of magnitude, enabling accurate inference and uncertainty quantification for large samples. Through extensive simulations, we demonstrate SINGER’s enhanced accuracy and robustness to model misspecification compared to existing methods. We illustrate the utility of SINGER by applying it to African populations within the 1000 Genomes Project, identifying signals of local adaptation and archaic introgression, as well as strong support of trans-species polymorphism and balancing selection in HLA regions.

## 1 Introduction

Many problems in genomics depend on computational methods to infer genealogical information from a large collection of DNA sequences and to interpret the reconstructed trees. In particular, genealogical approaches have been fundamental in understanding human genetic variation [28, 46, 58] and have laid the groundwork for numerous computational methods used in biomedical research. These methods include haplotype phasing when sequencing human genomes [3] and predicting statistical power in genome-wide association studies [29, 53]. In species with recombination, such as humans, the genealogical relationship cannot be represented by a single tree. Instead, millions of different trees exist across the genome, with each position in the genome typically having its own tree that only minimally differs from trees at nearby sites. The collection of all these trees, along with the set of recombination points that create new trees, is represented by the Ancestral Recombination Graph (ARG) and the corresponding generative model is called *the coalescent with recombination* [13, 21].

Although simulating under the coalescent with recombination is straightforward [2, 22, 27], inferring ARGs from genetic variation data remains a challenging problem. This challenge arises because the state space is extremely large, while the mutations that inform the genealogical relationship at any given position in the genome are limited. ARGs can be built iteratively by finding the branches and the times at which the *n*th lineage joins the partial ARG for the first *n −* 1 genomes, a process referred to as “threading.” By utilizing an approximation known as the sequentially Markov coalescent (SMC) [36, 37, 57] and formulating the threading problem as a hidden Markov Model (HMM), in combination with a clever Markov chain Monte Carlo (MCMC) method, ARGweaver [44] can sample genome-wide ARGs from the approximate posterior distribution for a moderate sample size. However, it comes with substantial computational overhead, rendering it impractical for more than a few tens of individuals.

Recently, significant advances have been made in scaling up ARG reconstruction to tens or hundreds of thousands of genomes. Tools like Relate [50] and tsinfer/tsdate [28, 58] utilize an efficient HMM by Li and Stephens [33] to infer a sequence of local tree topologies along the genome, followed by branch length estimation. ARG-Needle [60] constructs ARGs using the aforementioned threading approach, similar to ARGweaver, together with several heuristics and approximations to achieve scalability. The ability to infer genome-wide ARGs for large samples opens up new research directions [16, 17] and has enabled a series of ARG-based applications in population and statistical genetics research [9, 10, 14, 19, 24, 25, 41, 47, 51, 56].

Despite the progress mentioned above, there are significant limitations in current ARG inference methods, which impede ARG-based analyses of whole-genome sequencing (WGS) data. First, although recent methods are highly scalable, this increased speed comes at the cost of accuracy in key aspects of the reconstructed ARG, such as coalescence times [59] and recombination events [7]. For example, Relate and tsinfer+tsdate show notably reduced accuracy for ancient coalescence times, diminishing their effectiveness in applications involving ancient times, such as detecting balancing selection. Additionally, neither Relate nor tsinfer explicitly infers the branch position or the time of recombination events, and it is challenging to extract identity-by-descent (IBD) information from Relate’s internal data representation. Second, scalable methods generally do not explore alternative ARG topologies, often reconstructing only a single ARG topology. Some also overlook uncertainty in coalescence time estimates. As we demonstrate in this article, accurate sampling of ARG topologies can considerably enhance statistical inference, particularly for local gene tree analysis, where a point estimate can be quite noisy. Applications such as local ancestry inference, introgression detection, and selection analysis would be challenging without proper confidence intervals. For instance, CLUES [51], a likelihood-based method for inferring selection and allele frequency trajectories, benefits from incorporating a sample of local trees rather than relying solely on a point estimate. Third, most ARG reconstruction methods assume a simplistic prior, such as a constant-size panmictic population and the absence of selection, and are not robust against the violation of these assumptions. This limitation has consequential implication on applications, as many human populations have undergone complex demographic changes, including bottlenecks and recent expansions [32, 48]. Similarly, the influence of background selection is pervasive, profoundly shaping the diversity landscape [38, 40].

To address the issues mentioned above, we here introduce a new Bayesian ARG inference method, SINGER (**S**ampling and **IN**ferring of **GE**nealogies with **R**ecombination). It achieves all the functionalities of ARGweaver – including MCMC-based posterior sampling, exploration of topology space, tracking of recombination events, etc. – while being at least an order of magnitude faster. Through extensive simulations, we demonstrate that SINGER attains higher accuracy in several crucial aspects of ARG inference compared to competing methods. They also exhibit greater robustness to various common sources of model misspecification. We highlight the utility of our method by applying it to African populations within the 1000 Genomes Project, revealing signals of local adaptation and archaic introgression, as well as strong evidence of trans-species polymorphism and balancing selection in HLA regions.

## 2 Results

### 2.1 An overview of the SINGER algorithm

SINGER samples ARGs iteratively by adding one haplotype at a time via an operation called threading [44]. Conditioned on a partial ARG for the first *n−*1 haplotypes, the threading operation samples the points at which the lineage for the *n*th haplotype joins the partial ARG. SINGER solves this threading problem by first building an HMM with branches as hidden states and sampling a sequence of joining branches along the genome from the posterior, using stochastic traceback (Figure 1A). Then, SINGER builds another HMM with joining times as hidden states, conditioned on these sampled joining branches (Figure 1B). We refer to these two steps as “branch sampling” (Section 4.1) and “time sampling” (Section 4.2), respectively. We note that this two-step threading algorithm is approximative. However, compared to ARGweaver’s HMM, which treats every joining point in the tree as a hidden state, the two-step threading algorithm is much faster because the number of hidden states is substantially smaller.

**Figure 1:**
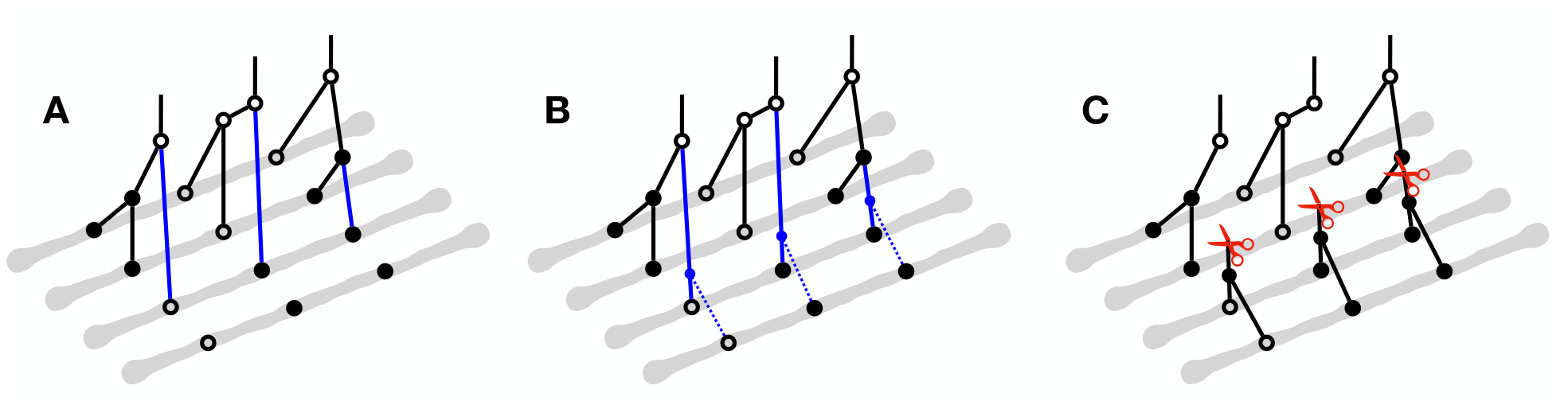
Method overview. The gray lines represent haplotypes, while the circles indicate the allelic states of nodes in coalescent trees. Hollow circles correspond to ancestral alleles and solid circles derived alleles. In panels A, B and C, a partial ARG for the first three haplotypes has already been constructed and a fourth haplotype is about to be threaded onto this partial ARG. (A) The initial step in threading the fourth haplotype involves sampling the joining branch (highlighted with blue) in each marginal coalescent tree of the partial ARG, a process we call “branch sampling”. (B) Following the determination of the joining branches, the next step is to sample the joining time for each of these joining branch. This step is referred to as “time sampling”. (C) To propose an update to an ARG in MCMC, we introduce cuts (illustrated by red scissors) to a sequence of marginal coalescent trees to prune subtrees, and then re-graft them by solving the threading problem for the sub-ARG above the cuts. This proposal is called “Sub-Graph Pruning and Re-grafting (SGPR)”.

Additionally, to explore the ARG space (both topology and branch lengths) according to the posterior distribution, we perform MCMC using a new proposal which we call Sub-Graph Pruning and Re-grafting (SGPR). Briefly, an SGPR operation first prunes a sub-graph by introducing a cut to a randomly chosen tree and then extends the cut leftwards and rightwards (Figure 1D); in Supplementary Section A.4, we show that the first step of this operation is equivalent to the removal step in the so-called “Kuhner move” [30, 35]. However, our SGPR operation differs substantially from the Kuhner move in the re-graft step; the latter samples from the prior by simulation, while our method utilizes the threading algorithm to sample from the posterior. The simulation procedure in the Kuhner move does not take data into account and hence is unlikely to reach a state with improved likelihood; in contrast, our threading algorithm encourages compatibility with data, which likely leads to better updates. Compared to previous methods, SGPR thus has a much better acceptance ratio while introducing large updates to the ARG (Supplementary Section A.4), resulting in better convergence rate and mixing of the MCMC.

Lastly, to adjust for some of the biases arising from the approximations in the algorithm, for every thinning interval of the MCMC algorithm, we rescale branch lengths (Section 4.3), using an algorithm we call “ARG re-scaling” (see Section 4.3 for details).

### 2.2 Performance benchmarks on simulate data

We first benchmarked the performance of various ARG inference methods using simulated data.

#### Setup

We utilized *msprime* [26] to simulate 1 Mb regions for sets of 50 or 300 haplotypes, using *μ* = *r* = 2 *×* 10^*−*8^, where *μ* and *r* respectively denote the per-generation per-base pair (bp) mutation rate and recombination rate. We compared the results of SINGER with those of popular ARG inference methods, namely ARGweaver, Relate, tsinfer+tsdate, and ARG-Needle. For Relate and tsinfer+tsdate, the results are based on posterior averages on the fixed topology estimated by these methods. ARG-Needle employs posterior averages of joining times when threading. SINGER and ARGweaver can average over the posterior of topologies, and our results for these methods are based on 100 sampled ARGs. To ensure a fair comparison, we used the same burn-in and thinning intervals across all methods. Detailed methodology can be found in Section 4.5.

#### Coalescence time accuracy

To assess the accuracy of inferred coalescence times, we compared the ground truth pairwise coalescence times and the inferred ones for randomly chosen 100 pairs of leaf nodes, as in YC Brandt et al. [59]. Pairwise coalescence times play an important role in the application of inferred ARGs, for example for demography inference [50], GWAS [10, 60], and in evolutionary studies [56]. For simulated data with 50 haplotypes, SINGER had higher accuracy than all other methods, with ARGweaver and Relate performing similarly, and tsinfer+tsdate performing the worst (Figure 2A). ARG-Needle only works for more than 300 sequences so it was not included in this comparison for 50 haplotypes. For 300 haplotypes, we compared only SINGER, Relate, tsinfer+tsdate and ARG-Needle, as the sample size is too large for ARGweaver. SINGER achieved the highest accuracy, while Relate and ARG-Needle performed similarly, with tsinfer+ts-date again performing the worst (Supplementary Figure S17). The improvement of SINGER compared to ARGweaver might be due to a better mixing efficiency in MCMC and a more flexible time discretization scheme (resulting in finer bins).

**Figure 2:**
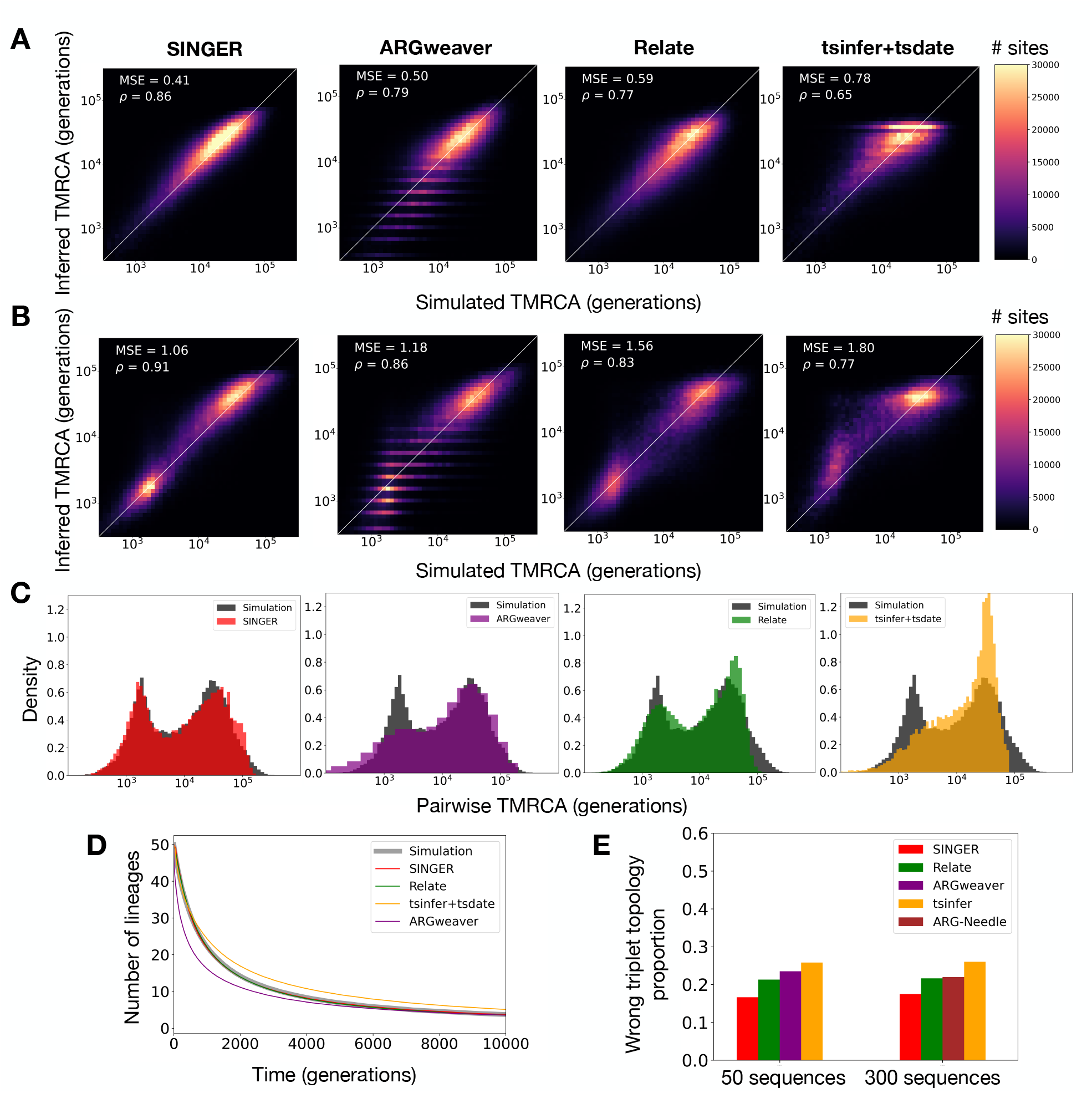
Performance benchmarks on coalescence time and topology inference. (A) Inferred pair-wise coalescence times compared with the ground truth in simulations involving 50 sequences under a constant population size scenario. (B) Similar to panel A but for data simulated under an inferred population size history for CEU. (C) Inferred distribution of pairwise coalescence times (colored) compared with the ground truth (black) from simulations under the same CEU demography as in panel B. (D) Genome-wide average of the number of lineages as a function of time for 50 sequences under a constant population size history, compared with the ground truth in simulations. (E) The proportion of triplet topologies that are incorrectly inferred for 50 and 300 sequences under a constant population size history. Due to runtime constraints, ARGweaver is not benchmarked for 300 sequences.

We also compared with the pairwise coalescent approach that performs inference for each pair of sequences separately, independently of other sequences. Specifically, we considered a recently proposed method, Gamma-SMC [49], which is ultra fast. Comparing Supplementary Figure S10 with Figure 2A suggests that SINGER can substantially outperform Gamma-SMC, whereas Relate and tsinfer+tsdate do not seem to improve on Gamma-SMC in terms of MSE or correlation.

Another benchmark we considered is the genome-wide average number of lineages as a function of time in marginal trees; this statistic is relevant to the inference of demography and selection. ARGweaver underestimated many recent coalescence times, resulting in the number of lineages dropping too fast compared to the expectation (Figure 2D). This is consistent with the results in Figure 2A, as well as the finding by Speidel et al. [50] that ARGweaver tends to underestimate times. We also found that tsdate tends to overestimate coalescence times substantially (Figure 2D). In contrast, we observed that the results of Relate and SINGER agree well with the expectation (Figure 2D).

#### Tree topology accuracy

To evaluate the accuracy of the inferred tree topologies, we employed the triplet distance, defined as the fraction of 3-leaved subtrees with different topologies in a given pair of trees. This metric is of particular interest because the accuracy of applications such as imputation and local ancestry inference (with two source populations) depends on the accuracy of triplet topologies. On average, SINGER achieved the lowest triplet distances to the ground truth (Figure 2E). Again, we observed that ARGweaver is not as accurate as SINGER. This reduced accuracy can be due to ARGweaver’s less efficient MCMC mixing and the presence of polytomies in the trees inferred by ARGweaver as a result of its time discretization.

#### Robustness to model misspecification

One advantage of our method is that they are more robust to model misspecification; specifically, it is less affected by using a wrong reference effective population size *N*_*e*_ or not accounting for population size changes. When we performed inference using an *N*_*e*_ off by a factor of 5, the coalescent times inferred by SINGER were less affected compared to those inferred by Relate and tsinfer+tsdate, which showed systematic downward biases (Supplementary Figure S11).

We also simulated data under an inferred population size history for CEU [42, 54], which contains a bottleneck and recent expansion. Performing ARG inference for these data showed that not only does SINGER more accurately infer the coalescence times than ARGweaver, Relate and tsinfer+tsdate (Figure 2B), but that it also accurately captures the bi-modality in the pairwise coalescence time distribution caused by the bottleneck (Figure 2C). Relate can incorporate population size changes, but it requires the user to run a separate module of re-estimating branch lengths and inferring coalescent rates themselves, which takes even longer than running Relate itself. In contrast, SINGER automatically adjusts branch lengths with ARG re-scaling and infer coalescence times accurately with almost no additional computational overhead, as the ARG re-scaling algorithm is very efficient. However, ARG-Needle requires a size history to be provided beforehand in order to adjust its coalescence times, and is not able to handle a unknown size history. We point out that a method that can jointly re-estimate branch length and population size history is likely to further improve the results of SINGER.

#### Runtime comparison

As described in Methods, the initialization and subsequent MCMC steps in SINGER and ARGweaver all utilize threading algorithms. Therefore, we compared the runtime of the threading operation as a function of the number of leaves in the partial ARG. Compared to ARGweaver, SINGER’s threading is around 10*×* faster. Moreover, as described in Section 2.2, the MCMC scheme in SINGER is substantially more efficient than ARGweaver’s, which implies that it requires many fewer MCMC iterations for mixing, thereby reducing the computational cost even further, resulting in a total speedup of *∼* 400*×*. We have also implemented automatic parallelization tools in SINGER to sample ARGs for multiple genomic regions simultaneously. In the real data application described below, performing 10,000 MCMC iterations for 200 African whole-genome sequences took about 2 days on Intel Xeon E5-2643 v3 Processors (3.4 Ghz) with 120 cores in total.

#### Accuracy of mutation and recombination inferences

We also benchmarked the estimation of allele ages using inferred ARGs. We omitted ARG-Needle and ARGweaver in this benchmark, since ARG-Needle does not map mutations to the inferred genealogies, and it is difficult to retrieve mutational mapping and ages from the output of ARGweaver. For simulated data with 50 sequences, SINGER noticeably outperformed Relate and tsinfer+tsdate in this task (Figure 3A). The results for 300 sequences are illustrated in Supplementary Figure S18, which suggests that SINGER remains more accurate than Relate and tsinfer+tsdate for larger sample sizes.

**Figure 3:**
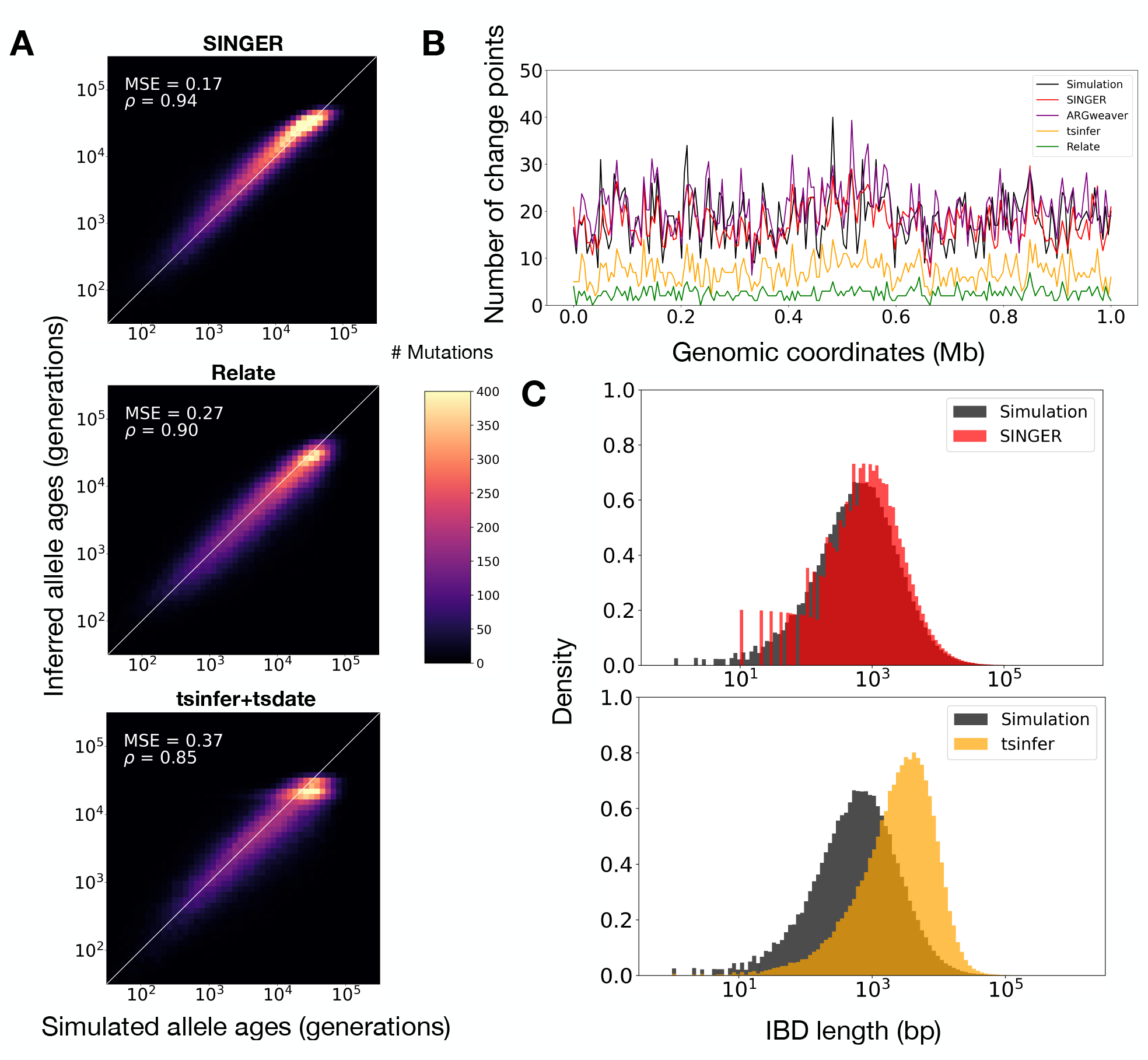
Benchmarks on mutation and recombination inference for data simulated with 50 sequences and constant population size. (A) Inferred allele ages compared with the ground truth. (B) Inferred number of recombination breakpoints in 5 kb genomic windows compared with the ground truth. (C) The length distribution of pairwise IBD in the inferred ARGs compared with the ground truth.

In another benchmark, we compared the number of inferred recombination breakpoints to the ground truth in each 5 kb window. Only ARGweaver and SINGER produced accurate estimates (Figure 3B). Both Relate and tsinfer missed many recombination events, a finding consistent with earlier studies [7].

Another way to assess the accuracy of recombination inferences is to examine the distribution of pairwise IBD length, as IBD segments are disrupted by recombinations. In this analysis, ARG-weaver and Relate were excluded; the former due to difficulties in extracting IBD information from its output, and the latter because it does not ensure node persistence among marginal trees. The results, illustrated in Figure 3C, show that SINGER accurately captured the distribution of pairwise IBD length, while tsinfer substantially overestimated the IBD length, aligning with previous findings [7].

#### Comparison of MCMC convergence

In humans and many other organisms, the genome-wide average of recombination rate is similar to the average mutation rate, which leads to substantial uncertainty in ARG inference. Therefore, it is important to obtain samples from the posterior distribution and ensure that uncertainties are well characterized, rather than relying solely on a point estimate. To assess MCMC convergence, we obtained 100 posterior MCMC samples from each of ARGweaver, Relate, and SINGER, using the same burn-in and thinning intervals for all methods.

To assess the effectiveness of sampling ARGs from the posterior distribution, we employed the same benchmark as in YC Brandt et al. [59]. This involved analyzing rank plots of pairwise coalescence times and quantifying deviations from the uniform distribution, which would be achieved by a perfect sampler from the posterior distribution [6, 52]. A rank plot is a histogram of the rank of a parameter sampled from the prior relative to the posterior sample. Ideally, a converged and well-mixed MCMC should produce ranks that follow a uniform distribution. In contrast, a U-shaped rank plot suggests sampling from an under-dispersed distribution [6, 52]. We observed that the rank plots for SINGER are much closer to the uniform distribution compared to those for ARGweaver and Relate (Figure 4B).

**Figure 4:**
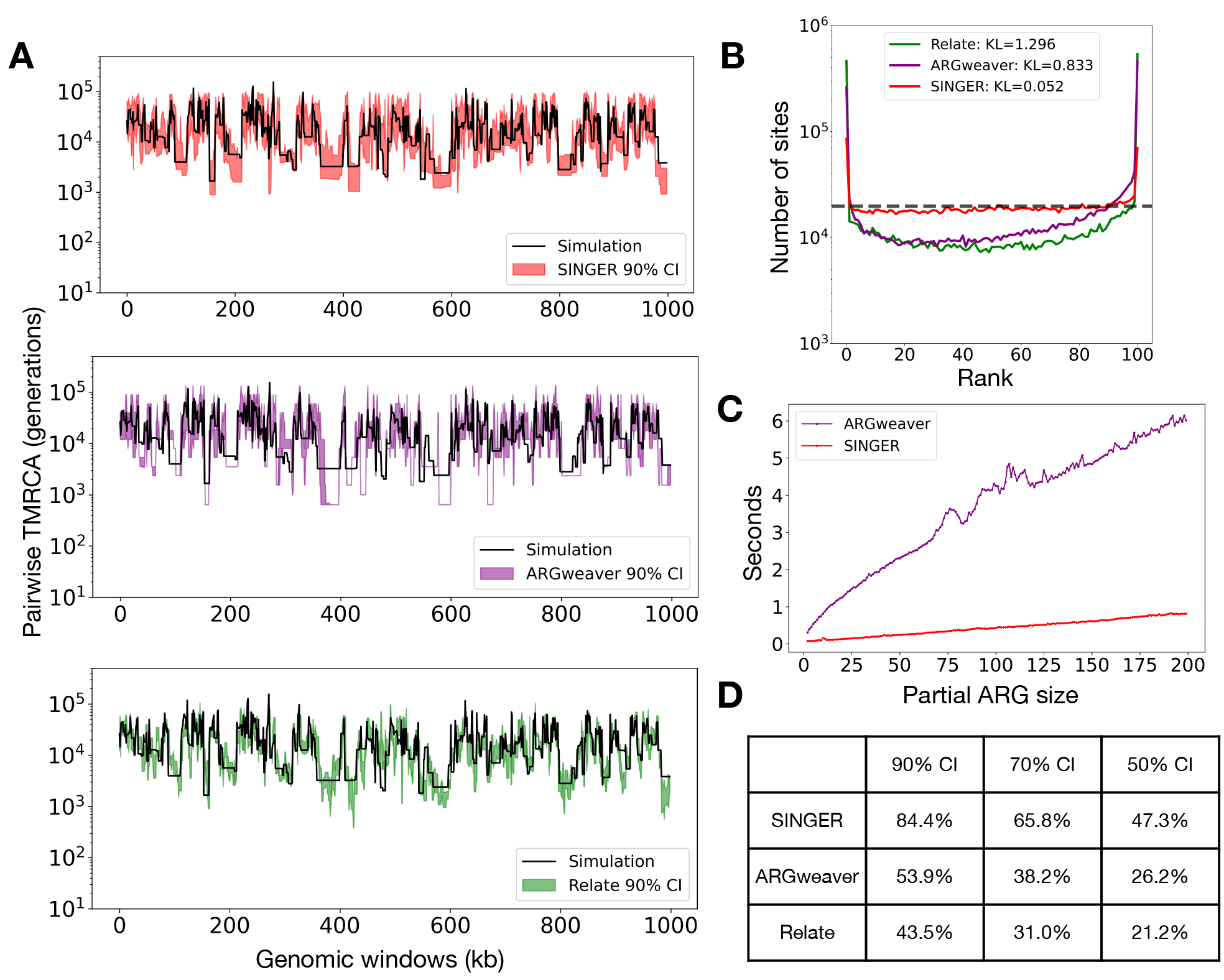
Properties of ARG samples and runtimes. (A) Empirical 90% credible intervals for pairwise coalescence times as inferred by SINGER, ARGweaver and Relate. (B) Rank plots of pairwise coalescence times in MCMC samples. A perfect sampler from the posterior distribution would achieve the flat dashed line, corresponding to the uniform distribution. The Kullback–Leibler (KL) divergence is utilized to quantify the deviation from the uniform distribution. (C) The runtime of the threading algorithm as a function of the partial ARG size (measured by the number of leaves), for ARGweaver and SINGER. (D) The empirical coverage of the ground truth pairwise coalescence time by the credible interval, for different nominal levels.

The rank plot is closely related to the coverage property of empirical credible intervals (CIs). For each position in the genome and for a given pair of haplotypes, we defined the empirical 90% credible interval by selecting the 5th to the 95th percentile of pairwise coalescence times from the sampled ARGs (Figure 4A). This approach was similarly applied to the 70% and 50% credible intervals. The 90% credible interval covered the ground truth in only 44% of instances for Relate and 54% for ARGweaver. In contrast, the coverage was substantially better for SINGER, at 85% (Figure 4C). CIs at other levels also showed superior coverage for SINGER (Figure 4C).

Furthermore, our benchmark showed that even with 40 times longer thinning intervals than SINGER, ARGweaver still slightly underperforms in the inference of pairwise coalescence time and coverage properties of CIs (Supplementary Figure S12). This underscores the advantage of the SGPR updates over ARGweaver’s MCMC algorithm. Furthermore, when combined with our faster threading algorithm, it implies that ARGweaver takes hundreds or even thousands times longer to reach performance levels comparable to those of SINGER. For Relate, even with a long thinning as suggested by the documentation, the CI coverage is still substantially lower than the nominal levels (Supplementary Section B.3). This is likely due to the fact that Relate only samples coalescence times with a fixed topology, in contrast to SINGER, which sample both topologies and coalescence times.

### 2.3 Applications to African WGS data from the 1000 Genomes Project

We applied SINGER to 200 whole-genome sequences from five African indigenous populations (GWD, YRI, ESN, LWK and MSL) in the 1000 Genomes Project [4], with 40 genomes drawn uniformly at random from each population. To demonstrate the utility of SINGER in population genetic analysis, we considered the inference of a few different types of evolutionary signals, including local adaptation, ancient balancing selection, and ancient introgression.

#### Diagnostics of the ARGs sampled by SINGER

We examined the sampled ARGs to check for convergence of the Markov chain to stationarity and to ensure that the ARGs used in our population genetic analyses are sampled past proper burn-in (Supplementary Section C.3). The chains generally showed good convergence to stationarity (Supplementary Figure S15). Additionally, we validated the accuracy of the sampled ARGs by comparing the inferred average pairwise coalescence times (scaled by 4*N*_*e*_*μ*) to the empirical average pairwise diversities (based on SNPs) in 500 kb windows, which generally showed high concordance. In contrast, we observed that the ARG reconstructed by tsinfer+tsdate [58] substantially under-estimated the genome-wide variation of diversity (Supplementary Figure S14). This is possibly because different genomic regions have different *N*_*e*_ values as a result of varying levels of background selection [5, 23]. As shown earlier in Section 2.2, SINGER is more robust to this type of model misspecification even if a fixed *N*_*e*_ is used in the algorithm.

#### Signatures of local adaptation

Local adaptation refers to population-specific selective sweeps, potentially due to selective pressures from special local environments. Genomic regions involved in local adaptation would exhibit reduced diversity for the specific populations experiencing selection, especially when compared to other populations without the sweep (Figure 5B). However, SNP-based inference of local diversity can be rather noisy at fine scales (Supplementary Figure S13). On the other hand, with branch length estimates from accurately inferred ARGs, fine-scale diversity can be estimated more accurately. As shown in Supplementary Section B.4 and Supplementary Figure S13, we observed that SINGER produces more accurate estimates of fine-scale diversity compared to Relate and tsinfer+tsdate. This provides a new opportunity to use ARG-based estimates of population-specific fine-scale diversity to study local adaptation. In particular, we note that this ARG-based approach can still be applied to detect a complete sweep with the beneficial allele fixed in the population, whereas methods like IHS [55] and the test proposed in Relate [50] that depend on segregating variants would suffer.

**Figure 5:**
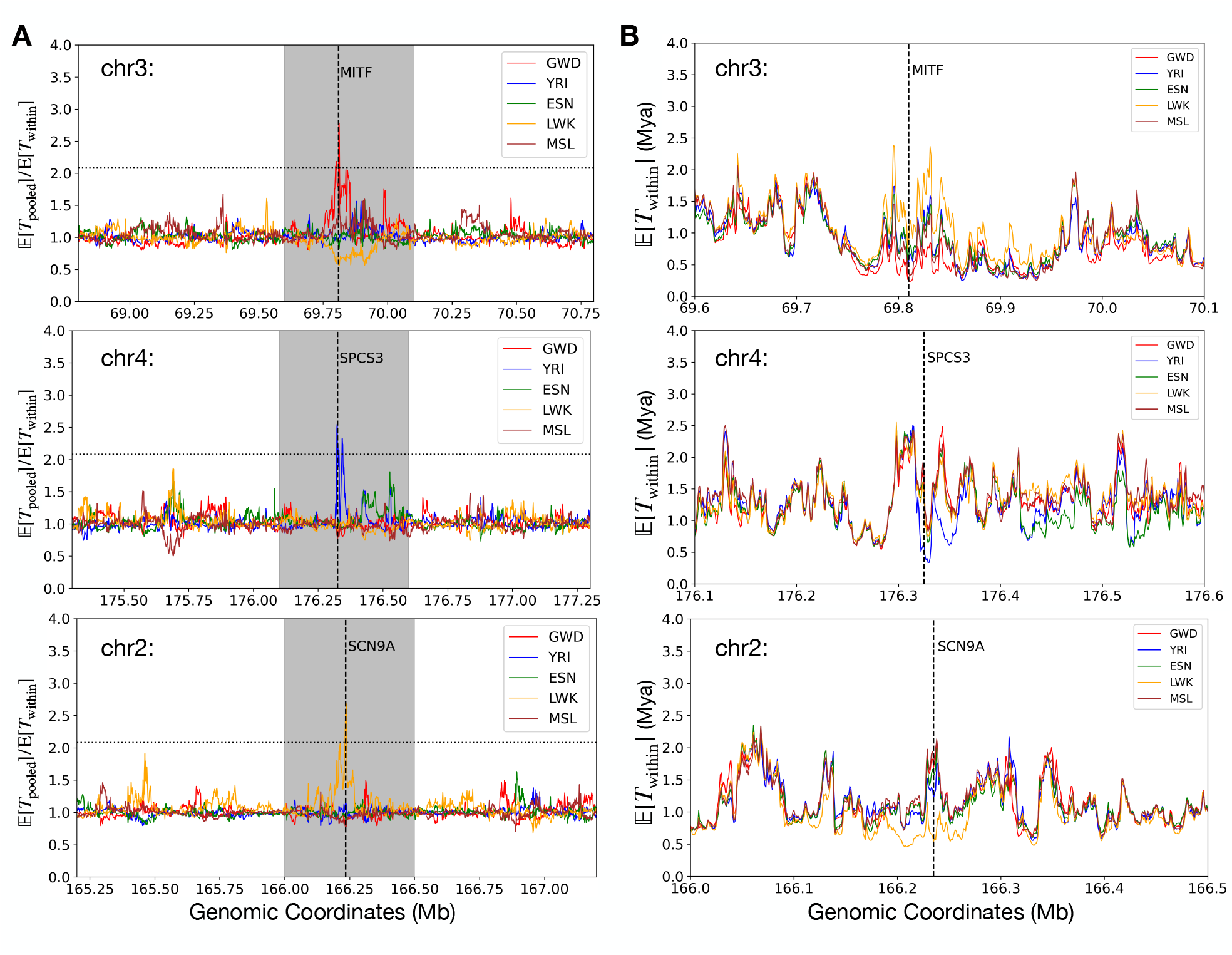
ARG-based detection of local adaptation in Africa. (A) The ratio of the average pair-wise coalescence time *T*_pooled_ in the pooled sample (combining all five populations) to the average population-specific pairwise coalescence time *T*_within_, for every 1 kb window. In each plot, the horizontal black dashed line denotes the genome-wide 99.99% quantile, and the gray shaded area corresponds to a 50 kb window surrounding the peak. A significant peak in this ratio may signal local adaptation in the corresponding population. The positions of these peaks are marked by vertical dashed lines, and the genes overlapping with these signals are indicated. (B) The average *T*_within_ for each population, zoomed into the gray regions highlighted in panel A.

To find population-specific reduction in local diversity, we partitioned the genome into non-overlapping 1 kb windows and then, for each window, computed the ratio of the ARG-based diversity estimate for the combined sample to the ARG-based population-specific diversity estimate for each of the five populations; reductions in local diversity would show up as peaks when these ratios are plotted along the genome. The full list of regions harboring elevated levels of the ratio for each population can be downloaded from the links provided in the Data Availability section. Analyzing the overlap with genic regions led to several interesting findings, a few of which we highlight in Figure 5. For example, we found that the gene *MITF* has experienced a putative sweep in GWD; this gene has been reported to be functionally related to skin pigmentation by encoding a melanocyte-inducing transcription factor [31]. Around *MITF*, we observed substantial differences in the ARG-based local diversity estimate across the five populations, consistent with previous findings concerning pigmentation variation within Africa [11]. In YRI, we found that *SPCS3* may have experienced a local sweep; this gene encodes an immune-related protein believed to impact virion production of flavivirues such as West Nile Virus and Yellow Fever virus [61]. This is concordant with the report of the spread of these diseases in Nigeria [1]. Lastly, we found that *SCN9A*, which encodes a voltage-gated sodium channel involved in the perception for pain [45], has substantially reduced diversity in LWK compared to other populations.

#### Balancing selection in the HLA locus

The Human Leukocyte Antigen (HLA) locus comprises a cluster of genes on human chromosome 6 which encode transmembrane proteins that present antigen peptides to T cells. This region is known to be the most diverse region in the human genome, and it has been hypothesized to be under extreme balancing selection to maintain its high diversity in order to handle various immune challenges [12, 34]. There has been evidence of trans-species polymorphism for some alleles across primates, which otherwise is very rare [12].

The ARGs inferred by SINGER show extremely ancient pairwise coalescence times in the HLA locus, with many regions harboring coalescence times older than the human-chimpanzee divergence time (Figure 6C). This result is consistent with the long-standing hypothesis of strong balancing selection in this locus and the observed patterns of trans-species polymorphisms. The human-chimpanzee divergence time has been debatable, with estimates ranging from 5-12 Mya [39]. While many genes in the HLA region do not show strong evidence of coalescence times older than the human-chimp split (e.g., *TAP1, TAP2*, and *TAPBP*), many do, including *HLA-A, HLA-DRB1*, and *HLA-DRB6*. Unsurprisingly, there is no noticeable differences across the five populations, as the polymorphism has been maintained since ancient times. In contrast, the ARG inferred by [58] using tsinfer+tsdate displays no such extreme coalescence times (Supplementary Figure S19), which might be due to the method’s lack of robustness to model misspecification in regions substantially deviating from the prior of selective neutrality.

**Figure 6:**
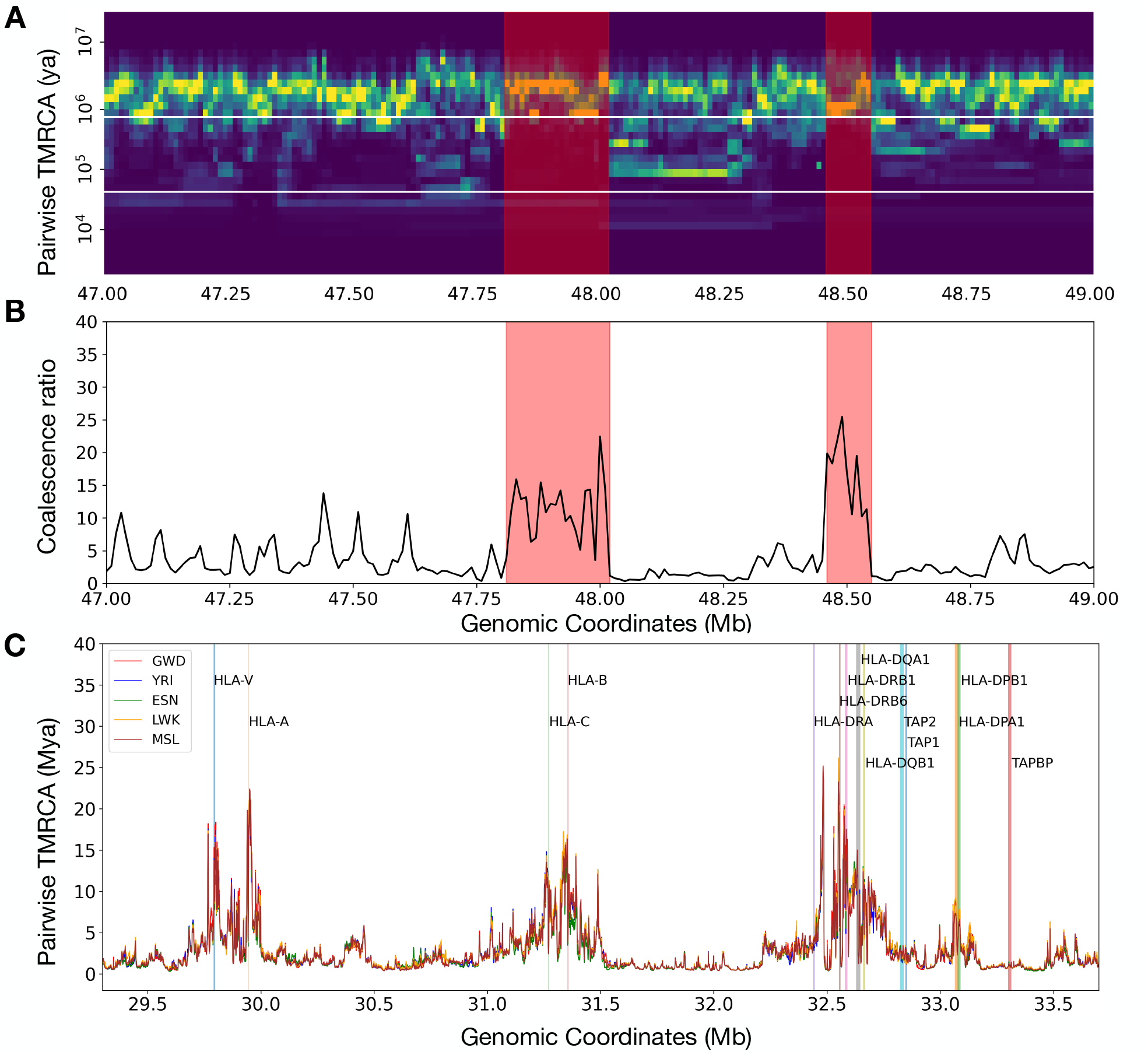
ARG-based detection of archaic introgression tracts and signatures of balancing selection. (A) Identification of potential archaic introgression tracts. For a given leaf node, its pairwise coalescence times with every other leaf node in the marginal tree are summarized as a distribution. In the plot, each column represents such a distribution from marginal trees within a 10 kb window. The two white horizontal lines delineate the interval between the introgression time and the split time. A tract indicative of introgression should exhibit a depletion of coalescence events within this interval and an enrichment of coalescence events above the split time. Regions shaded in red denote putative introgression tracts. (B) The ratio of pairwise coalescence density above the split time to that within the interval between the introgression time and the split time. (C) The average population-specific pairwise coalescence time *T*_within_ in the HLA locus. There are pronounced signals of trans-species polymorphism and balancing selection for several HLA genes.

#### Ancient introgression in Africa

It has been hypothesized that there may have been introgression from unknown “ghost” archaic hominins into ancient African individuals [15]. Identification of genomic tracts in modern African genomes resulting from such introgression is a challenging task, especially when there is no known genome of the source hominins. However, ARGs can facilitate this task by utilizing the following observation: for a genomic region with an introgressed tract in a given haplotype, coalescence between that haplotype and other haplotypes will be depleted in the interval between the introgression time and the split time of modern humans from the “ghost” population. This is similar to the “long branch” signals mentioned in [50], but expressed in the pairwise coalescence space.

However, long branches in ARGs can be rather sensitive to incorrect topology inference; specifically, the introgressed lineage can group incorrectly with the ancestral lineages of non-introgressed sequences, thereby destroying the long branch. To mitigate this issue, we provide a new technique for introgression analysis based on the coalescence distribution heatmap. For any given sequence, we plot the distribution of the time of pairwise coalescence with the remaining sequences in every 10 kb window, as illustrated in Figure 6A, where each column corresponds to a 10 kb window. We found that using a collection of ARGs sampled from the posterior instead of a single reconstructed ARG is helpful, as the coalescence distribution can be rather noisy in the latter (Supplementary Figure S16). Using ARG samples with different topologies helps to smooth the heatmap (Supplementary Figure S16). This visualization is similar to that proposed by Schweiger and Durbin [49].

To test whether a given segment arose from an ancient introgression event, we can look for a depletion of probability mass in the aforementioned interval and an enrichment of mass above the interval. This approach is more robust than explicitly relying on long branches because slight misgrouping would still lead to probabilistic depletion in the interval, whereas the long branch would be disrupted completely. Following Durvasula and Sankararaman [8], we used 43 kya and 625 kya for the introgression time and split time, respectively. If we plot the ratio of coalescence probability above the split time to that in the interval between introgression and split times, introgression tracts should appear as peaks, as illustrated in (Figure 6B), which shows two potential archaic introgression tracts of length 210 kb and 90 kb (Figure 6A).

## 3 Discussion

In this article, we introduced SINGER, a new Bayesian inference method designed for efficiently sampling Ancestral Recombination Graphs from the posterior distribution. These methods implement an improved MCMC algorithm to explore the ARG space, thereby enabling accurate uncertainty characterization in both coalescence times and ARG topologies. Our approach represents the first MCMC algorithm capable of scaling to at least hundreds of whole-genome sequences while performing full posterior sampling of both ARG branch lengths and topologies. Compared to ARGweaver, our approach benefits from both a faster threading algorithm and a more efficient exploration of the ARG space, leading to improved MCMC mixing. In estimating key population genetic quantities — such as coalescence times, topologies, recombination densities, and allele ages — SINGER compares favorably with existing ARG inference methods, including ARGweaver, Relate, tsinfer+tsdate, and ARG-Needle. As we have demonstrated in our benchmarks, utilizing posterior samples can noticeably enhance inference accuracy and effectively quantify estimation uncertainty. Last but not least, SINGER exhibits greater robustness to common sources of model misspecification, such as population size changes and background selection, without the necessity of explicitly modeling them or subsequent re-estimation and re-fitting.

We applied SINGER to the WGS data of African individuals from the 1000 Genomes Project, uncovering various signals of selection and archaic introgression. Utilizing population-specific fine-scale diversity estimates derived from our inferred ARGs, we identified genomic regions showing evidence of local adaptation. In addition, we found strong evidence of balancing selection and trans-species polymorphisms in the HLA region, and mapped genes associated with the peaks in these signals. Lastly, we employed a visualization technique, the coalescence distribution heatmap, to identify genomic regions consistent with a specific model of archaic introgression.

We note that our proposed approach for detecting archaic introgressed tracts requires a demographic model of introgression. However, there is ongoing debate regarding the timing, strength, and even the existence of archaic introgression into African populations, as highlighted in Ragsdale et al. [43]. In this regard, the tracts identified by our approach will be reliable only if the assumed times of introgression and population splits are reasonably accurate. Moreover, the detection of introgressed tracts using sampled ARGs warrants further methodological development. Our proposed heatmap of coalescence distribution summarizes the genealogical information within a collection of sampled ARGs, providing a basis for developing a systematic algorithm. We defer to future research the challenges of selecting an appropriate critical value for the coalescence ratio indicative of introgression, as well as determining the boundaries of introgressed tracts.

There are a few limitations of our method, along with potential strategies to address them. First, although SINGER is substantially faster and more scalable than other methods capable of full posterior sampling of ARGs, real data applications often require a large number of MCMC iterations to effectively infer the posterior distribution. To reduce this computational load, we are developing a method to sample ARG branch lengths given a fixed ARG topology, which could then be incorporated into the current MCMC algorithm of SINGER. Additionally, MCMC convergence tends to be slower for large samples, so there is a need to devise more efficient ARG exploration strategies.

Second, while SINGER shows improved robustness to model misspecification compared to other ARG inference methods, inferring population size history and re-estimating ARG branch lengths, similar to the approach used by Relate [50], may enhance accuracy further. We note that Relate’s algorithm for inferring population size history and re-estimating branch lengths cannot be readily applied to other ARG inference methods, because it relies on the special data structures of Relate. On the other hand, tsdate’s algorithm for sampling ARG branch lengths works for the data structure of SINGER, but it only supports constant population size. More generally, in line with ARGweaver-D Hubisz et al. [20], SINGER could potentially be extended to include more complex demographic models.

Third, for partitioning the genome into bins, SINGER currently uses equal-sized bins for programming simplicity, but it would be better to use dynamic bin sizes according to the inferred recombination density. Dynamic binning can be especially important for regions with recombination hotspots, since we allow at most one recombination between adjacent bins and so it would be less likely to underestimate recombinations by introducing more bins. Moreover, using larger bins for regions with low recombination rates will reduce the runtime. The recombination landscape in the human genome has punctuated hotspots separated by lower recombination rate regions, so the dynamic binning will likely lead to substantial overall computational savings.

Lastly, our method requires phased genomes as input and assumes accurate phasing. Unfortunately, obtaining high-quality phased data is often challenging, especially for understudied population and non-model organisms. Therefore, enhancing ARG inference methods to support partially or completely unphased data would increase their utility. Additionally, considering that all data sets contain some degree of phasing errors, the ability to automatically correct poorly phased heterozygote sites based on sampled ARGs would be advantageous. This feature would be particularly important in joint analyses of ancient and modern genomes, as ancient samples are often poorly phased or unphased.

## 4 Methods

### 4.1 Branch sampling

To speed up the computation, we first partition the genome into equal-sized bins, where the bin size is chosen to be 4 *×* 10^*−*3^*/*4*N*_*e*_*r*. We then construct an HMM indexed by these bins (loci) to sample the branches at which the lineage for the *n*th haplotype joins the partial ARG for the first *n −* 1 haplotypes. The state space *S*_*𝓁*_ for locus *𝓁* comprises all the branches in the marginal tree for locus *𝓁* in the partial ARG and some branches from earlier loci. The precise definition of the state space can be found in Supplementary Section A.1. For locus *𝓁*, we use *B*_*𝓁*_ to denote the branch onto which the lineage for the *n*th haplotype joins. If the partial ARG does not already contain a recombination event between loci *𝓁 −* 1 and *𝓁*, then the transition probability of the HMM is defined as

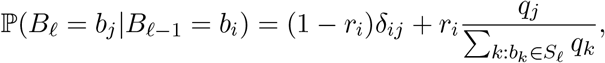

where *b*_*i*_ *∈ S*_*𝓁−*1_, *b*_*j*_ *∈ S*_*𝓁*_, and *r*_*i*_ denotes the branch-specific recombination probability for branch *b*_*i*_. The definition of *r*_*i*_, *q*_*i*_ can be found in Supplementary Section A.1.5. The structure of this transition probability is similar to that of the Li-Stephens model [33], but with branch-specific recombination and re-joining probabilities. This allows us to reduce the HMM computational complexity to be linear with respect to the number of hidden states, as in the Li-Stephens model.

We restrict the final ARG to have at most one recombination event between adjacent loci. Therefore, if the partial ARG already contains a recombination event between loci *𝓁 −* 1 and *𝓁*, the threading operation is not allowed to introduce an additional recombination between these two loci and the transition probability in this case is defined similarly to that in Rasmussen et al. [44]; the details are provided in Supplementary Section A.1.4 and Supplementary Figure S3.

### 4.2 Time sampling

Conditioned on a sequence of joining branches along the genome resulting from the branch sampling algorithm, the time sampling step proceeds in a similar fashion as in PSMC [32] but with the restriction that, for each genomic locus *𝓁*, the coalescence time should reside between the two endpoints of the joining branch for locus *𝓁*. To accelerate the computation, we implemented the linearization technique of Harris et al. [18]. The details are provided in Supplementary Section A.2.

### 4.3 ARG re-scaling

Given an inferred ARG, we partition the time axis into non-overlapping windows such that the total branch length across all marginal trees (weighted by the span of each tree) in each time window is the same; by default 100 windows are chosen. We then count the number of mutations falling into each of these windows. If a mutation falls on a branch striding multiple windows, then its contribution to the mutation count for each window is given by the proportion of the branch that overlaps with the window. We rescale each window size such that the expected number of mutations for the window matches the empirical count. This is essentially a window-specific re-scaling to better match the mutation clock. Further details can be found in Supplementary Section A.3. ARG re-scaling is performed after the initialization and every thinning step.

### 4.4 Sub-graph pruning and re-grafting (SGPR)

In the MCMC algorithm, we propose updates to the current ARG by first removing some branches following a cut and then re-grafting from the breakpoint.

To remove a branch from a given marginal tree, we make a random cut on the tree; the probability that a given branch gets cut is proportional to its length. We can extend the cut leftwards and rightwards along the genome, removing the partial branch from the cut to its upper endpoint of the branch. Typically the cut will not extend over the entire chromosome; rather, the extension width will be the same as the span of the ancestral segment corresponding to the branch that got cut. The details can be found in Supplementary Section A.4 and Supplementary Figure S8.

To re-graft the branch from the breakpoint, we use the same threading algorithm as previously described, with the only difference being that now we only consider the sub-ARG above the break-point. We show in Supplementary Section A.4.3 that, assuming the threading algorithm samples approximately from the posterior, the acceptance probability is typically much higher than that of previous proposals [30, 44], thereby improving convergence and mixing in MCMC.

### 4.5 Simulation and benchmarking details

All coalescent simulations in this article were carried out using *msprime* [26] with *r* = *m* = 2 *×* 10^*−*8^ and *N*_*e*_ = 1 *×* 10^4^. We simulated 50 datasets each with 50 sequences over a 1 Mb region and 10 datasets each with 300 sequences over 1 Mb. For simulations with 50 sequences, we also simulated with CEU population size history https://github.com/PalamaraLab/ASMC_data/tree/main/demographies estimated from SMC++ [54]. We ran all inference methods with these true parameter values.

As for MCMC sampling, we uniformized the number of iterations and thinning scheme across all methods. We drew 100 samples with the thinning interval set to 20 for ARGweaver, Relate, and SINGER, and used 1,000 iterations for burn-in.

For all simulation benchmarks involving ARGweaver and SINGER, posterior averages were taken for the statistics of interest, such as pairwise coalescence time, allele age, etc. Since Relate outputs averages of MCMC iterations while tsdate results are averages from a probability table, we simply use their results from a single output as they are effectively posterior averages.

### 4.6 Running SINGER on African WGS data

We used 200 whole genomes from 5 African indigenous populations (GWD, YRI, ESN, LWK, and MSL) in the 1000 Genomes Project, with 40 genomes drawn uniformly at random from each population. The per-generation per-bp mutation rate (*μ*) and recombination rate (*r*) were both set to 1.2 *×* 10^*−*8^ in SINGER. In order to determine the effective population size *N*_*e*_, we matched the empirical average pairwise difference (*π ≈* 0.001) with the theoretical expectation (4*N*_*e*_*μ*), which led to a choice of *N*_*e*_ = 20, 000. We ran SINGER for 10,000 iterations, with the first 4,000 iterations as burn-in. We then took 100 samples from the rest, thinning every 60 iterations.

## Data availability

SINGER source code can be downloaded from https://github.com/popgenmethods/SINGER. We have uploaded the inferred ARG samples (100 samples) and gene targets for local adaptation to Zenedo at https://doi.org/10.5281/zenodo.10437053, https://doi.org/10.5281/zenodo.10467284, and https://doi.org/10.5281/zenodo.10467509.

## Acknowledgements

We thank Matthew Rasmussen and Melissa Hubisz for helpful correspondence regarding ARG-weaver; Vince Buffalo for discussion on background selection; Debora Brandt for discussion on the benchmark pipeline; Joshua Schraiber for advice on data analysis; Montgomery Slatkin and Alyssa Fortier for discussion on HLA; James Santangelo, Chuck Langley, Nathaniel Pope, and Jules Perez for testing the software. This research is supported in part by an NIH grant R56-HG013117.

## Notes

### Competing Interest Statement

The authors have declared no competing interest.

