## Supplementary Information for "Robust and Accurate Bayesian Inference of Genome-Wide Genealogies for Large Samples"

March 16, 2024

#### Contents

A Method details 3
A.1 Branch sampling 3
A.1.1 Representative joining time and joining probability for branches 3
A.1.2 Deterministic approximation 4
A.1.3 Emission probability 4
A.1.4 Transitions induced by an existing recombination in the partial ARG 5
A.1.5 Transitions with a new recombination 8
A.2 Time sampling 9
A.2.1 Transition and emission probabilities 9
A.2.2 Linearization of the forward algorithm for type A transitions 10
A.2.3 Type B and type C transitions 11
A.2.4 Inference of recombination times 12
A.3 ARG rescaling 12
A.4 Sub-Graph Pruning and Regrafting (SGPR) 13
A.4.1 How to prune a sub-graph from an ARG 13
A.4.2 Comparison of SGPR and the Kuhner move 14
A.4.3 How to regraft the sub-graph to generate an updated ARG 16

B Simulation benchmarks 17
B.1 Comparison with PSMC 17
B.2 Robustness to model misspecification 18
B.3 The impact of thinning on rank plots 19
B.4 Fine-scale diversity estimation 20

**C Applications to African data from the 1000 Genomes Project** **20**

**D Additional Supplementary Figures** **24**

### A Method details

The key components of SINGER include branch sampling, time sampling, ARG re-scaling, and Sub-Graph Pruning and Re-grafting (SGPR). Here we describe these procedures in detail. Figure S1 illustrates the terminology we employ.

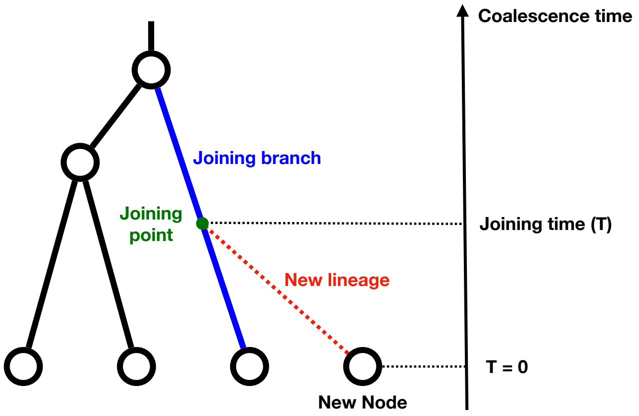

Figure S1: Illustration of the terminology used. “New node” refers to the node being threaded onto the partial tree, while “new lineage” refers to the branch connecting the new node to the partial tree. “Joining branch” refers to the branch in the partial tree that the new lineage attaches to, and “joining point” refers to the location of attachment. The time of the joining point is called “joining time”.

#### A.1 Branch sampling

In the branch sampling step, we construct an HMM with branches as hidden states to “thread” a new node onto a sequence of marginal trees. Let $S_\ell$ denote the state space for locus $\ell$. There are two key quantities we need to compute for each branch in the marginal tree: (1) A representative joining time for the branch, which determines the probability of seeing mutations on the new lineage; and (2) the probability of joining the branch under the coalescent process, which serves as the model prior and is needed for calculating transition probabilities in the HMM (Section A.1.5). These are detailed below.

##### A.1.1 Representative joining time and joining probability for branches

For threading a new node onto a single marginal tree $\Psi$, let $T$ be the joining time of the new node and $\lambda_\Psi(t)$ the number of lineages at time $t$ in the tree. The exceedance probability $\bar{F}_\Psi(t) := \mathbb{P}_\Psi(T > t)$ and the density function $f_\Psi(t)$ of $T$ are given by

$$\bar{F}_\Psi(t) = \exp\left( - \int_0^t \lambda_\Psi(x) dx \right), \tag{1}$$

$$f_\Psi(t) = -\frac{d}{dt}\bar{F}_\Psi(t) = \lambda_\Psi(t)\bar{F}_\Psi(t). \tag{2}$$

The probability of coalescing onto a branch $b_i$ that spans the time interval $[x, y]$ can be calculated as

$$p_i = \int_x^y \frac{f_\Psi(t)}{\lambda_\Psi(t)} dt = \int_x^y \bar{F}_\Psi(t) dt.$$

We choose a representative joining time $\tau_i$ for each branch $b_i$ for the calculation of the transition (Section A.1.5) and emission probabilities (Section A.1.3), as detailed in the next section.

##### A.1.2 Deterministic approximation

Suppose the partial ARG has $n$ leaf nodes. For $n \gg 1$, $\lambda_\Psi(t)$ is almost deterministic and is well approximated by its expectation (Frost and Volz, 2010)

$$\lambda_\Psi(t) \approx \frac{n}{n + (1-n)\exp(-\frac{t}{2})}.$$

Using this result, we can approximate $\bar{F}_\Psi(t)$, $f_\Psi(t)$, and $p_i$ as

$$\bar{F}_\Psi(t) = \exp\left(-\int_0^t \lambda_\Psi(x)dx\right) \approx \frac{\exp(-t)}{[n + (1-n)\exp(-\frac{t}{2})]^2}$$

$$f_\Psi(t) = \lambda_\Psi(t)\bar{F}_\Psi(t) \approx \frac{n\exp(-t)}{[n + (1-n)\exp(-\frac{t}{2})]^3}$$

$$p_i = \int_x^y \bar{F}_\Psi(t)dt \approx \left\{\frac{-2}{(1-n)^2}\log\left[n + (1-n)\exp(-t/2)\right] - \frac{2n}{(1-n)^2}\frac{1}{n + (1-n)\exp(-t/2)}\right\}\Bigg|_x^y.$$

We choose the representative joining time $\tau_i$ for branch $b_i$ using a heuristic:

$$\lambda(\tau_i) = \sqrt{\lambda(x)\lambda(y)}$$

$$\tau_i = \lambda^{-1}(\sqrt{\lambda(x)\lambda(y)}),$$

where

$$\lambda^{-1}(l) = -2\log\left(\frac{n - nl}{l - nl}\right)$$

is the inverse function of $\lambda(\cdot)$. This choice works well empirically.

We note that $p_i$'s do not necessarily sum to 1, but they will be used in a "scaled-by-sum" fashion in transition probabilities (Section A.1.5). We apply the above deterministic approximations to all marginal trees in the partial ARG, thereby achieving massive computational savings in branch sampling.

##### A.1.3 Emission probability

We assume that mutations arise on each branch according to a Poisson point process with intensity $\theta/2$, independently of all other branches. Here, $\theta = 4N_e\mu$, where $N_e$ denotes the reference effective population size and $\mu$ the per-bp, per-generation mutation rate. When the new node attaches to the joining branch, it will create a new lineage and bisect the joining branch into two (Figure S2); these three branches are involved in the calculation of emission probability. We first impute the '0/1' state at the joining point under parsimony, and calculate the mutation probability for each of the 3 branches. The product of these 3 probabilities is then defined as the per-bp emission probability (Figure S2), and we multiply these probabilities over the base pairs in each locus to obtain the per-locus emission probability.

A special case is when the new lineage joins the branch above the root. Specifically, the allelic state at the new root is set to 0 with probability $p_{\text{root}}$ or 1 with probability $1 - p_{\text{root}}$, By default, we use $p_{\text{root}} = 0.5$ for un-polarized data; it can be set to a high value (e.g., 0.99) if the data are

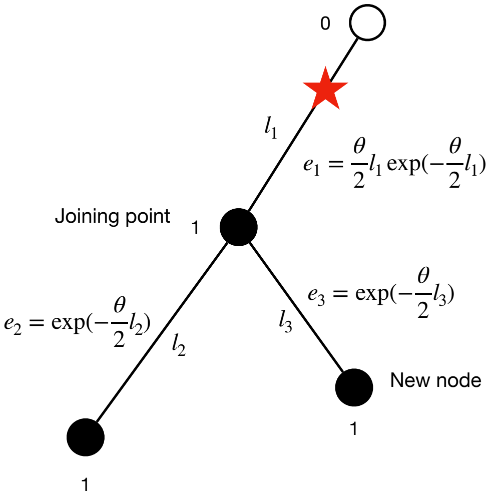

Figure S2: The emission probability calculation. We first impute the '0/1' character state at the joining point, which in this case should be 1, so a mutation (red star) has to be placed on the upper half of the joining branch. We can calculate the probability of each of three branches, and the product of them will be the final emission probability.

polarized, where 0 denotes the ancestral state. We do not make any assumptions on the allelic state of other nodes of the tree. Our recommendations on how to set this parameter can be found at the GitHub documentation: `https://github.com/popgenmethods/SINGER`.

##### A.1.4 Transitions induced by an existing recombination in the partial ARG

When the partial ARG already contains a recombination event between a given pair of adjacent loci, we forbid the new lineage from introducing any new recombination event between those loci. However, the joining branch for the new lineage can switch between such loci simply by "hitchhiking" with the existing recombing lineage, as detailed below.

If we trace the joining branch in the adjacent marginal trees before and after a recombination, the correspondence is as shown in Figure S3, which is the same as in Rasmussen et al. (2014). Joining a segment with a certain color in the previous tree will necessarily mean joining the segment of the same color in the next tree, and vice versa. For example, if the new node joins the red branch in the first tree, then it has to join the red segment in the second tree; because of the recombination, the upper node of the red branch in the first tree no longer exists in the second tree and the red segment is only a portion of a longer branch in the second tree.

Hence, after a recombination in the partial ARG, some joining branches will be only partial branch segments instead of full branches (e.g., the red and blue segments in the second tree in Figure S3), and these partial segments arise from full branches of a previous tree but extended to the current position. We refer to these segments as "partial branches". We note that partial branches can go through multiple recombinations along the genome, during which they will become more and more fragmented. The state space for a given site consists of all full branches of the marginal tree at the site and a set of partial branches arising from previous sites. We do not keep track of every partial branch state, but only keep a partial branch state if it has a forward probability (in the forward algorithm of the HMM) larger than a threshold $\epsilon$ (usually set at 1%). In other words, we prune unlikely partial branches while running the forward algorithm at the same time. This controls the state-space size to be at most $2n - 1 + \frac{1}{\epsilon}$. Empirically, we observed that the

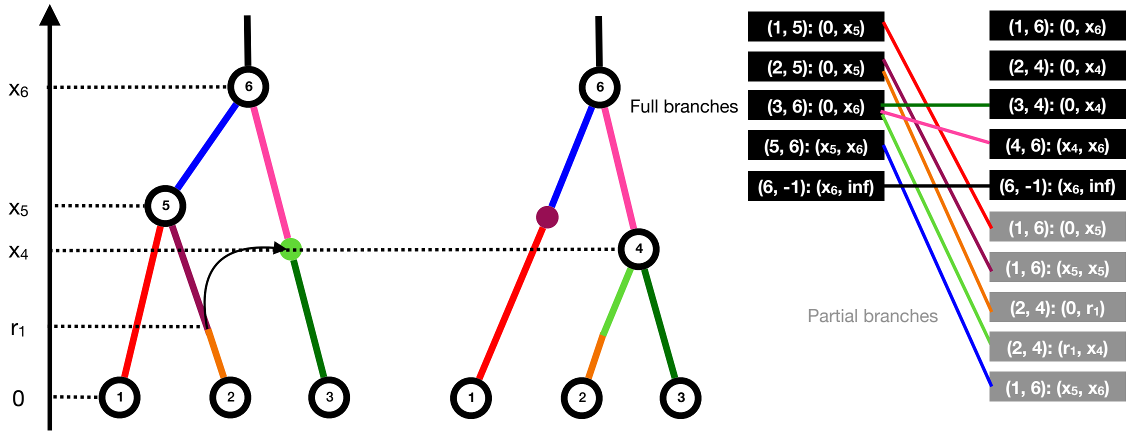

Figure S3: Branch correspondence (left) and HMM state spaces (right) before and after a recombination. When there is an existing recombination in the partial ARG, the joining point will shift between segments with the same color. This means if the new node joins the (partial) branch with one color in the previous tree, it must join the (partial) branch with the same color in the next tree, vice versa. On the right, we mark the state transitions with the same color of these segments correspondence. "$-1$" denotes a point at infinity. Note that a full branch in the previous tree might become a partial branch after the recombination (e.g., the red and blue), a full branch could also become 2 full branches (e.g., pink and dark green) and a branch could also be not affected at all (e.g. black root branch). A state is represented with the branch and the time interval on the branch, and all possible transitions are plotted (right). For example, the red segments correspond to $(1,5):(0,x_5) \to (1,6):(0,x_5)$ and purple segments correspond to $(2,5):(0,x_5) \to (1,6):(x_5,x_5)$.

state space size is only slightly larger than $2n-1$.

When a branch becomes multiple segments (e.g., in Figure S3, the full branch state $(2,5):(0,x_5)$ from the left tree breaks up into $(1,6):(x_5,x_5)$ and $(2,4):(0,r_1)$ in the right tree), the transition probability from the branch will be distributed to the segments proportional to the coalescence probability (obtained by integrating $\frac{f_\Psi(t)}{\lambda_\Psi(t)}$ over their respective time intervals in the branch). A more subtle case in Figure S3 is the full branch state $(3,6):(0,x_6)$ in the left tree which breaks up into three segments. Joining anywhere on the light green segment of the right tree will look like joining the light green dot on the left, so we set the time interval for the light green dot to be the time interval of the recombination arrow.

**Why not just use full branch states?** Here we explain why using only full branch states can be problematic. Consider the partial ARG shown in Figure S4A, which has three marginal trees spanning one locus each. Suppose a third sequence is being threaded onto it. If we constructed an HMM using only full branches, the state space for each locus would be as shown in Figure S4B. Since the partial ARG already contains recombination events between adjacent loci, the new lineage cannot introduce additional recombinations. In threading the third sequence, the transition $(1,5) \to (3,-1)$ is possible if the new lineage joins the branch $(1,5)$ above the recombination time $r_1$ in the first locus (Figure S4D). The transition $(3,-1) \to (1,4)$ is also possible if the new lineage joins the branch $(1,5)$ below $r_1$ in the first locus (Figure S4D). However, the joint move $(1,5) \to (3,-1) \to (1,4)$ is impossible in an ARG without introducing an additional recombination in the new lineage. This is effectively a consequence of decoupling the sampling of topologies and the sampling of coalescence/recombination times. Even under standard Markovian approximations to the ARG

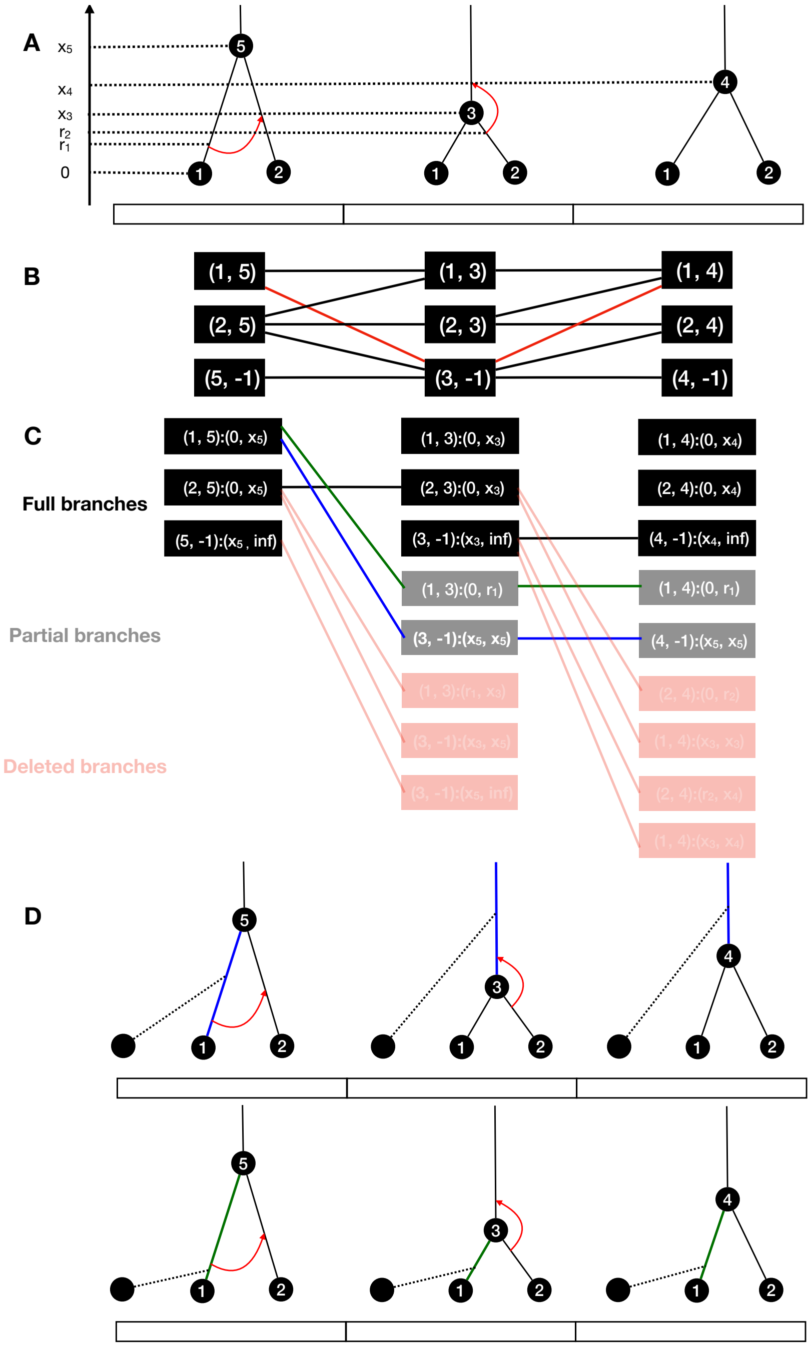

Figure S4: An example of the state space in the HMM and why only using full branches as hidden states will not work. Each hidden state consists of a branch and a time interval, and depending on whether the time interval covers the entire branch, a state can be either a full or partial branch, colored black or grey, respectively. The solid lines connecting states between adjacent columns indicate state transitions when no new recombination from the threaded node happens. In this example, because we do not allow more than one recombination to happen between adjacent loci, there should be no more new recombinations. (A) The partial ARG when threading the third leaf node; (B) What the HMM looks like when only using the full branches as hidden states, and the red path is allowed by the HMM but disallowed in an ARG; (C) The HMM with both full and partial branch states. Two valid path are marked with blue and green respectively; (D) The two different valid threadings which correspond to the HMM path in (C).

generation process, the Markovian property only holds exactly when recombination and coalescence times are represented in the state space. However, we can address this issue by introducing partial branch states.

Our proposed HMM is shown in Figure S4C. Every full branch in a marginal tree must become a hidden state, and the tricky part is birth and death of partial branch states. When transitioning from the first to the second locus, if the state for the first locus is $(1, 5) : (0, x_5)$, which is a full branch state, it becomes the partial branch state $(1, 3) : (0, r_1)$ or $(3, -1) : (x_5, x_5)$, depending on whether the joining point is lower or higher than the recombination time, respectively. We note that not all partial branch states are kept, as they must have forward probabilities larger than a threshold. Those below the threshold will be dropped, indicated by the red states in Figure S4C. In this example, both blue and green paths are valid in the HMM (Figure S4C), and they correspond to the two different threading operations shown in Figure S4D.

##### A.1.5 Transitions with a new recombination

Consider a pair of adjacent loci $\ell - 1$ and $\ell$, and let $b_i$ be the joining branch for locus $\ell - 1$ with representative time $\tau_i$. If the partial ARG does not already contain a recombination event between these adjacent loci, then the new lineage can undergo recombination with probability

$$r_i = 1 - \exp\Big( - \frac{\rho}{2}\tau_i \Big),$$

where $\rho = 4N_e rm$, with $r$ being the per-bp, per-generation recombination probability and $m$ the locus size. The transition probability of the HMM can be defined as

$$\mathbb{P}_\rho(B_\ell = b_j | B_{\ell-1} = b_i) = (1 - r_i)\delta_{ij} + r_i \frac{q_j}{\sum_{k:b_k \in S_t} q_k},$$

for $b_i \in S_{\ell-1}$ and $b_j \in S_\ell$, where

$$q_j = \begin{cases} r_j p_j, & \text{if state } b_j \text{ is a full branch,} \\ 0, & \text{if state } b_j \text{ is a partial branch.} \end{cases}$$

This has the interpretation that with probability $1 - r_i$ the new lineage will stay on the same joining branch due to the absence of recombination, whereas if a recombination happens to the new lineage, it will join branch $b_j$ proportional to $q_j$. The $q_j$ terms for partial branches are always 0, because after a recombination, the new lineage always joins a full branch at locus $\ell$. This also means that a partial branch state can be entered only when a full joining branch state from an earlier locus results in a partial branch state due to a recombination in the partial ARG.

In the sequentially Markov coalescent (SMC), given that a recombination happens in the new lineage, the conditional transition probability from $b_i$ to $b_j$ will depend on both $b_i$ and $b_j$, but to reduce computational complexity we drop the dependence on the previous joining branch and assume that the re-joining probability distribution is the same regardless of the previous joining branch. However, the choice of the re-joining probability distribution still needs to guarantee that the stationary distribution is $\mathbb{P}(B_\ell = b_i) = p_i$, in the most simple case when there is only one marginal tree in the ARG. This leads to choosing the re-joining probability distribution to be $\{q_j / \sum_{k:b_k \in S_\ell} q_k\}$. This also reflects the fact that lower branches are down-weighted when considering the re-coalescence after a recombination, because re-coalescence needs to be more ancient than the recombination event, which forbids joining branches lower than the recombination breakpoint.

#### A.2 Time sampling

Conditioned on a sequence of joining branches along the genome obtained from branch sampling, the joining times on these branches can be sampled using a modified PSMC (Li and Durbin, 2011) model with the state space for each locus restricted to the time interval of the sampled branch.

We note that although the state space in the branch sampling HMM involve both partial and full branches, only the sequence of full joining branches are used in time sampling step. Partial branches are only used when calculating the transition probability in certain cases (detailed in Section A.2.3 for type B)

This model, which is also an HMM, is detailed below.

##### A.2.1 Transition and emission probabilities

First, consider the simple case where the joining branch spans the entire non-negative half-line $[0, \infty)$. In this case, conditioned on there being a recombination in the new lineage, the probability density of the joining time transitioning from $s$ to $t$ is given by

$$q_0(t|s) = \int_0^{s \wedge t} \frac{1}{s} e^{-(t-u)} du = \begin{cases} \frac{1}{s}[1 - e^{-t}], & t < s, \\ \frac{1}{s}[e^{-(t-s)} - e^{-t}], & t \geq s. \end{cases}$$

Similarly, if we do not condition on there being a recombination,

$$q_\rho(t|s) = \begin{cases} \frac{1-e^{-\rho s}}{s}[1 - e^{-t}], & t < s, \\ e^{-\rho s}, & t = s, \\ \frac{1-e^{-\rho s}}{s}[e^{-(t-s)} - e^{-t}], & t \geq s, \end{cases}$$

and the corresponding cumulative distribution function is

$$Q_\rho(t|s) = \int_0^t q_\rho(x|s) dx = \begin{cases} \frac{1-e^{-\rho s}}{s}[t + e^{-t} - 1], & t < s, \\ \frac{1-e^{-\rho s}}{s}[s - e^{-(t-s)} + e^{-t}] + e^{-\rho s}, & t \geq s. \end{cases} \quad (3)$$

Now, suppose the joining branch at locus $\ell$ spans the time interval $[x_\ell, y_\ell)$. We partition this time interval into $d$ sub-intervals $[t_{\ell,0}, t_{\ell,1}), [t_{\ell,1}, t_{\ell,2}) \ldots, [t_{\ell,d-1}, t_{\ell,d})$, where $t_{\ell,0} = x_\ell$ and $t_{\ell,d} = y_\ell$, uniformly according to the exponential distribution with rate 1 (the default is partitioning every 5% quantile). In the time sampling HMM, these sub-intervals correspond to the states for locus $\ell$. For each sub-interval $[t_{\ell,i}, t_{\ell,i+1})$, we define the representative time $\tau_{\ell,i}$ as

$$\exp(-\tau_{\ell,i}) = \frac{\exp(-t_{\ell,i}) + \exp(-t_{\ell,i+1})}{2}.$$

Then, we define the transition probability from a sub-interval $[t_{\ell-1,i}, t_{\ell-1,i+1}) \subset [x_{\ell-1}, y_{\ell-1})$ at locus $\ell - 1$ to $[t_{\ell,j}, t_{\ell,j+1}) \subset [x_\ell, y_\ell)$ at locus $\ell$ as

$$q_{i,j}^{\ell-1,\ell} = \frac{Q_\rho(t_{\ell,j+1}|\tau_{\ell-1,i}) - Q_\rho(t_{\ell,j}|\tau_{\ell-1,i})}{Q_\rho(y_\ell|\tau_{\ell-1,i}) - Q_\rho(x_\ell|\tau_{\ell-1,i})}. \quad (4)$$

We define the emission probability in the same way as in branch sampling, depicted in Figure S2.

The state spaces for two consecutive loci in the time sampling HMM can be different if the joining branch changes between those loci. More precisely, there are three cases to be considered, as illustrated in Figure S5: (A) Neither the joining branch nor the partial ARG changes. (B) The

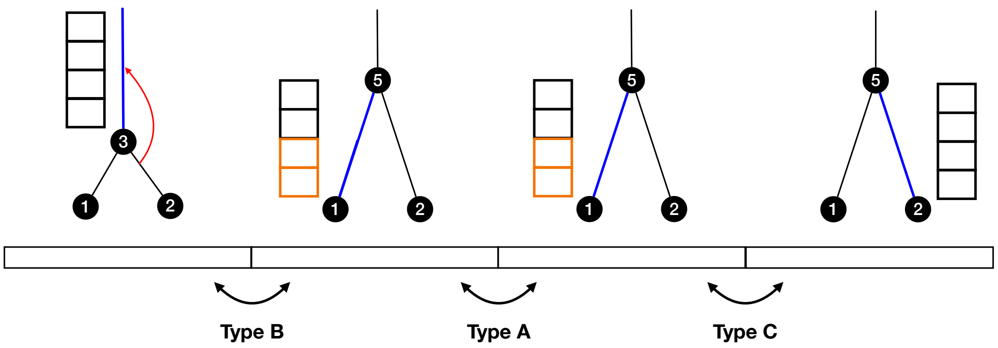

Figure S5: An illustration of the state space in the time sampling step. The blue branches denote a sequence of joining branches in different bins. The transition from the first to the second bin corresponds to type B, where the joining branch changes with partial ARG changing as well (recombination hitchhiking). The orange states are created to fill up the branch which is not reached by "recombination hitchhiking"; the transition from the second to the third bin corresponds to type A, where neither the joining branch nor the partial ARG changes. The state space will not change with transitions of type A; the transition from the third to fourth bin corresponds to type C, where the joining branches changes without partial ARG changing (new recombination).

partial ARG changes and the joining branch might change (recombination hitchhiking). (C) The partial ARG does not change but the joining branch does (new recombination). We note that both type B and type C transitions will involve either a previous recombination in the partial ARG or a new recombination in threading, but as the number of loci is typically much larger than that of recombinations, the majority of transitions will be of type A.

##### A.2.2 Linearization of the forward algorithm for type A transitions

The computational complexity for the general HMM forward algorithms is quadratic in the state space size, but, in the case of type A transitions, symmetry structures of the transition matrix in our HMM allow us to reduce the runtime to linear complexity (Harris et al., 2014). We note, however, that time sampling is not the computational bottleneck of SINGER. Because we sample from the posterior using the HMM with stochastic traceback, we only need to implement the forward algorithm.

Here, we follow the arguments in Palamara et al. (2018). For type A transitions between loci $\ell - 1$ and $\ell$, note that the state space is the same for the two loci; specifically, $t_{\ell-1,i} = t_{\ell,i}$ for all $i = 0, \ldots, d$. Hence, for ease of notation, we drop the dependence on $\ell - 1$ and $\ell$ in what follows.

First, note that (3) and (4) imply

$$q_{i,j} = q_{j+1,j}, \quad \text{for all } i > j, \tag{5}$$

whereas

$$\kappa_j := \frac{q_{i,j}}{q_{i,j-1}} = \frac{\exp(-t_j) - \exp(-t_{j+1})}{\exp(-t_{j-1}) - \exp(-t_j)}, \quad \text{for all } i < j. \tag{6}$$

(The right hand side of (6) does not depend on $i$, and we call it $\kappa_j$.) Let $\alpha_i(\ell)$ denote the forward probability at locus $\ell$ and state $i$, which corresponds to the joint probability of observed pairwise data for the first $\ell$ loci and the hidden state at locus $\ell$ being $i$. Let $e_j(\ell + 1)$ denote the emission probability at locus $\ell + 1$ from hidden state $j$. Then, the forward algorithm can be written as follows:

$$\begin{aligned}
\alpha_j(\ell+1) &= \sum_i \alpha_i(\ell) q_{i,j} e_j(\ell+1) \\
&= e_j(\ell+1)\left[\sum_{i<j} \alpha_i(\ell) q_{i,j} + \alpha_j(\ell) q_{j,j} + \sum_{i>j} \alpha_i(\ell) q_{i,j}\right] \\
&= e_j(\ell+1)\left[\sum_{i<j} \alpha_i(\ell) q_{i,j} + \alpha_j(\ell) q_{j,j} + \sum_{i>j} \alpha_i(\ell) q_{j+1,j}\right] \\
&= e_j(\ell+1)[S_j + \alpha_j(\ell) q_{j,j} + A_j q_{j+1,j}],
\end{aligned}$$

where $S_j := \sum_{i<j} \alpha_i(\ell) q_{i,j}$, $A_j := \sum_{i>j} \alpha_i(\ell)$, and the third equality follows from (5). Note that $S_j$ and $A_j$ for all $j = 0, \ldots, d-1$ can be computed recursively in linear time using

$$\begin{aligned}
S_j &= \alpha_{j-1}(\ell) q_{j-1,j} + \kappa_j S_{j-1}, \\
A_j &= \alpha_{j+1}(\ell) + A_{j+1},
\end{aligned}$$

with boundary conditions $S_0 = 0$ and $A_{d-1} = 0$, where the recursion for $S_j$ follows from (6).

##### A.2.3 Type B and type C transitions

**Type B transitions.** Here, the background partial ARG changes, whereas there may or may not be a change in the joining branch (e.g., the first transition in Figure S5). If the joining branch does not change, then the joining time also does not change in the time sampling HMM.

As we allow at most one recombination between any pair of adjacent loci, the joining branch can change in type B transitions only by "hitchhiking" an existing recombination in the partial ARG as described in Section A.1.4, in a way consistent with the choice of joining branches from the branch sampling process. In this case, each time sub-interval for the joining branch before the recombination is treated as a state in the time sampling HMM, and its corresponding state after the recombination will be constructed only if it is on the sampled joining branch. For example, in Figure S5, the first transition is type B, and the upper two time sub-intervals of the joining branch for the first locus will be on the branch $(5, -1)$ at the second locus after the transition, which is not consistent with the joining branch $(1, 5)$ sampled from the branch sampling process. Hence, these upper two time sub-intervals will not contribute to the calculation of new forward probabilities. Transition probabilities to the upper two sub-intervals on $(1, 5)$ at the second locus are defined in the same way as in Section A.1.4 for partial branches. Finally, when these "hitchhiked" states from before the recombination do not cover the entire branch, we introduce more sub-intervals to fill in the remainder, in the same fashion as in Section A.2.1. In Figure S5, transition probabilities from the first locus to these newly filled states (in orange) at the second locus are defined to be 0, but transition probabilities from the second locus to the orange states at the third locus are non-zero.

**Type C transitions.** If the joining branch changes between loci $\ell$ and $\ell+1$ due to a recombination in the new lineage (the third transition in Figure S5), we first build a new state space for locus $\ell + 1$, consisting of time sub-intervals of the new joining branch as described in Section A.2.1. Let $[t_{\ell,0}, t_{\ell,1}), \ldots, [t_{\ell,d_\ell-1}, t_{\ell,d_\ell})$ and $[t_{\ell+1,0}, t_{\ell+1,1}), \ldots, [t_{\ell+1,d_{\ell+1}-1}, t_{\ell+1,d_{\ell+1}})$ respectively denote the state spaces at loci $\ell$ and $\ell + 1$. Here, since we are already conditioning on having a new recombination (from the branch sampling process), the transition probability can be calculated by setting $\rho = \infty$ in (4). We update the forward probabilities using the standard recursion, as it is not a computational bottleneck in the time sampling HMM:

$$\alpha_j(\ell+1) = \sum_i \alpha_i(\ell) q_{i,j} e_j(\ell+1).$$

##### A.2.4 Inference of recombination times

Although the threading algorithm infers recombination events, it does not infer the exact timing of the recombination breakpoint on the recombining branch.

The recombination time is when the new lineage before and after the recombination get decoupled. The new lineage after recombination will wait from recombination time $x$ until joining time $v$ to re-coalesce with the new joining branch, which we call "re-coalescence event".

To sample the time of the recombination breakpoint, we note that under the SMC, given that the recombination breakpoint occurs at time $x$, the probability density of the re-coalescence event being at time $v$ is

$$p(x) = e^{-(v-x)}, \tag{7}$$

for $l < x < u$, where $l$ denotes the lower node age of the recombining branch, and $u$ the minimum of the joining times before and after the recombination. For the recombination time, we choose the median according to (7), conditioned on $l < x < u$.

#### A.3 ARG rescaling

Given an ARG, we partition the time axis into $J$ intervals $[t_0 = 0, t_1), [t_1, t_2), \ldots, [t_{J-1}, t_J = t_{max})$, where $t_{max}$ is the maximum node age before rescaling, such that in each interval $[t_i, t_{i+1})$ the ARG length is $\frac{1}{J}$ of total ARG length (the default value of $J$ is 100 in our implementation). Here, by "ARG length in an interval", we mean the sum of total branch length overlapping the interval from all marginal trees, weighted by their tree span. Let $m_i$ denote the number of mutations in the interval $[t_{i-1}, t_i)$. We note that some mutations may be mapped to a branch which spans more than one interval, in which case we assign fractions to each interval proportional to the overlap length.

If the total ARG length is $L(G)$, then the expected number of mutations in each time interval should be $\frac{\theta L(G)}{2J}$, so the scaling factor for the interval $[t_{i-1}, t_i)$ needed to match the observation with the expectation should be

$$c_i = \frac{2Jm_i}{\theta L(G)}.$$

We recursively scale and shift the intervals so that $[t_{i-1}, t_i)$ maps to $[\tilde{t}_{i-1}, \tilde{t}_i)$, where $\tilde{t}_0 := 0$ and

$$\tilde{t}_i = c_i(t_i - t_{i-1}) + \tilde{t}_{i-1},$$

for $i = 1, \ldots, J-1$. Then, a coalescence time $t \in [t_{i-1}, t_i)$ is rescaled to

$$\tilde{t} = c_i(t - t_{i-1}) + \tilde{t}_{i-1}.$$

Figure S6 illustrates this procedure for the case of a single coalescent tree.

This procedure is similar in spirit to the ARG normalization step in Zhang et al. (2023), as they also post-process the node ages in the ARG to improve time estimates. However, their ARG normalization assumes a known demography to generate the node age distribution by simulation and performs quantile matching based on that information. In contrast, our ARG rescaling strategy is self-contained and only utilizes information from the inferred ages of mutations.

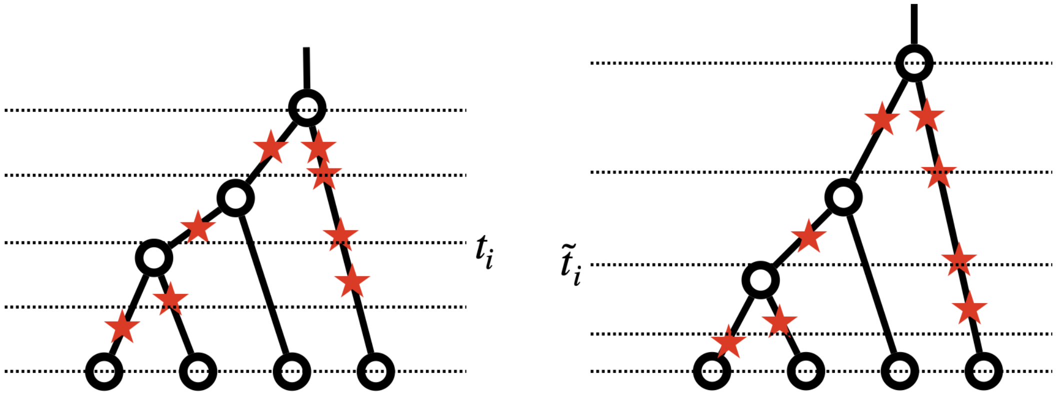

Figure S6: ARG rescaling from original ARG (left) to a new ARG (right), for simplicity we only show a single tree. We partition the ARG and count the number of mutations in each interval, and re-scale the interval length so that the expected number of mutations match the observed mutation counts. For example, the highest interval has more mutations mapped to it than the lowest interval, so it is widened and the latter is narrowed.

#### A.4 Sub-Graph Pruning and Regrafting (SGPR)

SGPR can be viewed as an extension of the Subtree Pruning and Regrafting (SPR) operation. SPR is a commonly used technique for modifying trees to explore the tree space in phylogenetics, by cutting a subtree and reconnecting it to another location in the remaining tree. In a similar vein, SGPR prunes a sub-graph from an ARG and reconnects it to another part of the remaining graph.

##### A.4.1 How to prune a sub-graph from an ARG

To prune a sub-graph from an ARG, we first introduce a random cut in a certain marginal tree. The cut usually extends to other trees in the flanking regions. To find the spatial span of the cut, we look for the equivalent point extending leftward and rightward in the tree sequence, where the rule for equivalency is illustrated in Figure S7. A cut located on a colored segment should always correspond to the segment with the same color in the other tree, except when the cut reaches a black dashed line, at which point the extension terminates. For a pair of adjacent loci with a recombination event occurring between them, the black dashed line (Figure S7) in each marginal tree corresponds to the part of the new lineage above the recombination breakpoint; these partial branches get decoupled by the recombination and equivalence of points cannot be established between them (leftwards or rightwards), and hence the extension of the cut has to terminate. For each marginal tree in the span of the cut, we remove between the cut and the upper node of the branch containing the cut (Figure S8).

A cut may or may not span the entire chromosome, and we keep a record of the rightmost position of the span to choose the next cut. If the rightmost position is the end of the chromosome, we start over from the beginning. Otherwise, the next cut will be chosen at the marginal tree at the rightmost position of the previous cut, according to the following steps:

1. Sample a cut time $t$ uniformly at random between 0 and the tree height (tree TMRCA).

2. Determine the branches intercepting time $t$.

3. Choose one of them uniformly at random and cut the chosen branch at time $t$.

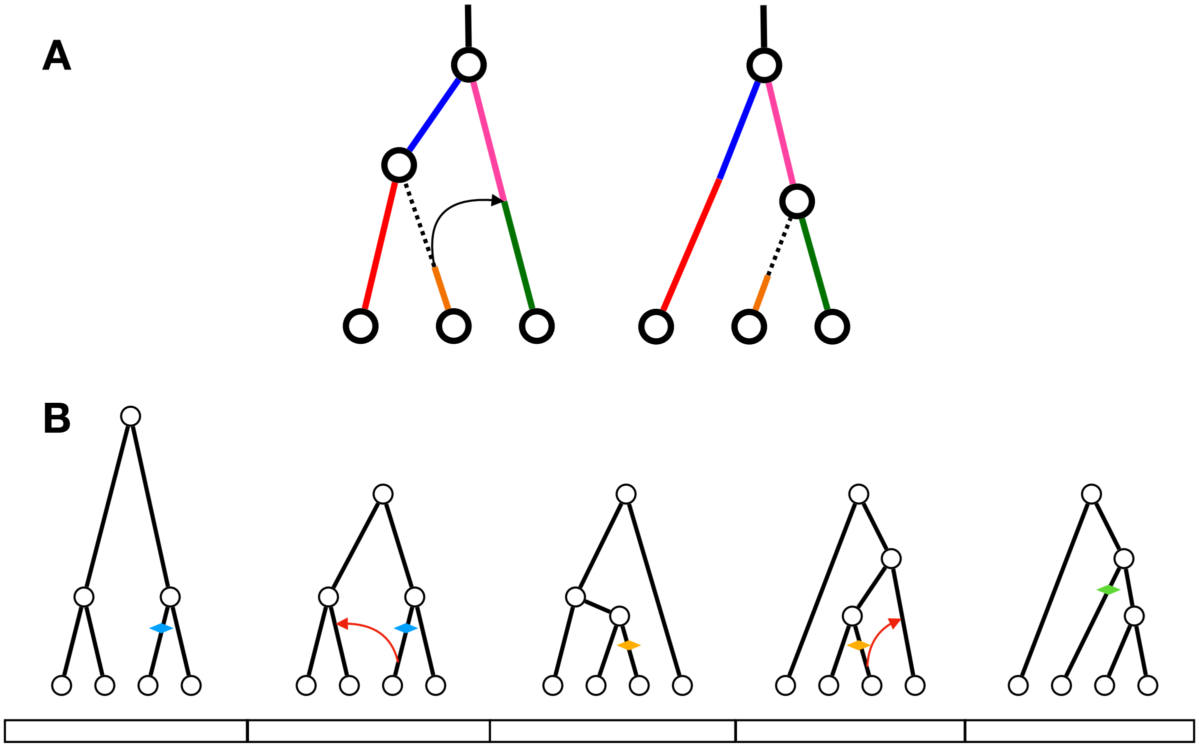

Figure S7: The correspondence of cut in adjacent trees and an example of cut extension. (A) If a cut is on the segment with a particular color in the first tree, then the cut will be on the segment with the same color in the second tree (at the same time), vice versa. If the cut is on the dashed segment, then it cannot be extended leftwards or rightwards. (B) In this example, we start by introducing a blue cut to the marginal tree in the first bin, and it extends to and terminates at the second bin. After SGPR induced by the blue cut, a new orange cut is introduced in th marginal tree of third bin, which spans over the third and fourth bin. A green cut in the fifth bin follows the SGPR induced by the orange cut.

##### A.4.2 Comparison of SGPR and the Kuhner move

Here we point out the similarities and differences between the Kuhner move (Kuhner et al., 2000) and our proposals (SGPR). The Kuhner move works with the temporal formulation of the ARG, and starts by picking a branch in the graph (e.g., the green branch in Figure S8A). Then, all history (meaning that all coalescence events and recombination events) for the ancestral material corresponding to this branch get removed from the graph (Figure S8B). Since we can translate the temporal representation of the ARG into a spatial representation comprising a sequence of marginal trees, the ancestry removal operation in the Kuhner move leads to a sequence of trees with portions of branches removed (Figure S8B). In other words, one can think of it as introducing a cut to a marginal tree and propagating the cut leftwards and rightwards. For every marginal tree affected, the part of the branch above the cut is removed (Figure S8B). This removal operation of the Kuhner move is the same as in SGPR.

The main difference is in the regraft step (Figure S8), in that the Kuhner move does this by simulating the coalescent with recombination process from the prior distribution, starting from the

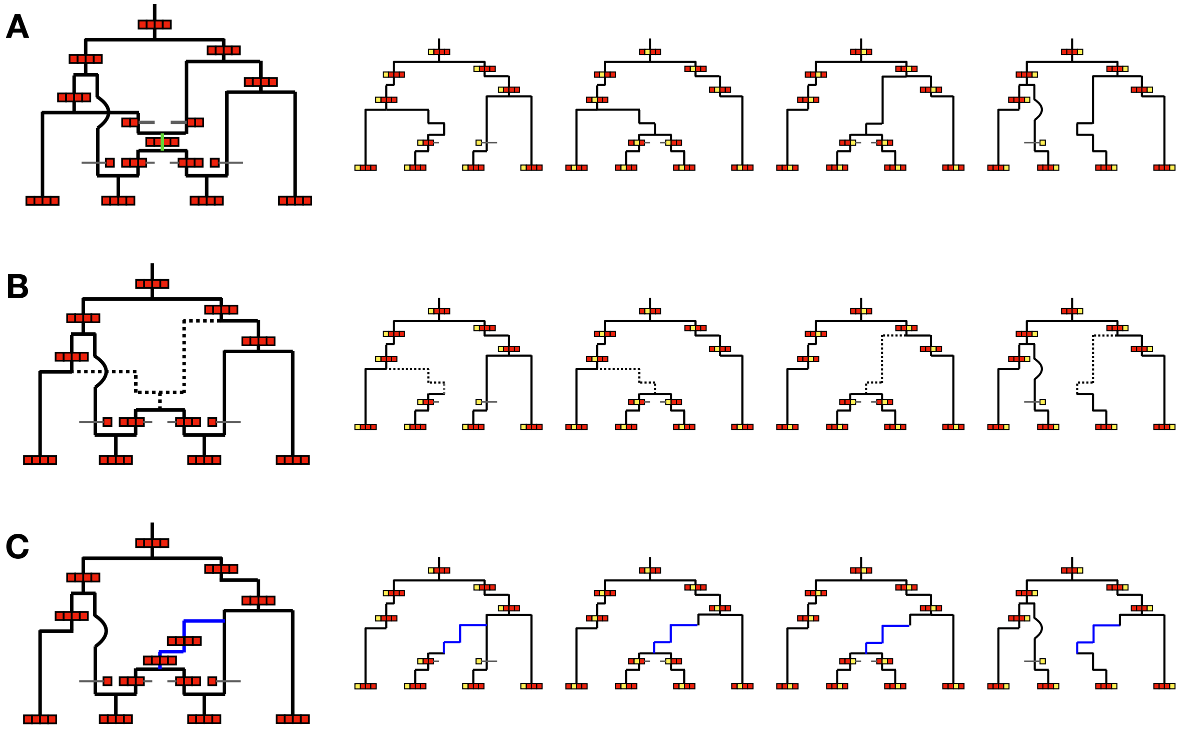

Figure S8: The connection between the Kuhner move and SGPR. ARG can be represented as the temporal network (left) or the spatial series of trees (right). (A) is the ARG before update in temporal network and spatial trees. In (B), the Kuhner move first picks a random branch (shown in green) in the network and removes any recombination or coalescence above from that branch, which is marked with dashed line (left). It is equivalent to choose a cut and trace it to the flanking region in SGPR (right). The partial branches connecting the cut to its upper node in every marginal tree is removed. In (C), the sub-graph has been re-grafted back with a new coalescent history, leading to potential changes in topology and branch length. For example, the recombination separating the second and third tree in the previous ARG (A) is gone.

cut. This is intuitively problematic because the regrafted genealogy receives no information from the data and might not be consistent with the data. This problem can be seen from the acceptance ratio of the Kuhner move (Mahmoudi et al., 2022):

$$A(G \to G') = \min\left\{1, \frac{B(G)\mathbb{P}(G'|D)}{B(G')\mathbb{P}(G|D)}\right\},$$

where $B(G)$ is the number of branches in the graph and $\mathbb{P}(G|D)$ is the likelihood of graph $G$ given the data $D$. This result implies that unless the likelihood of the newly proposed graph $G'$ is substantially better than before, the proposal will be unlikely to be accepted. However, this is very hard to achieve if the prior distribution is employed to complete the ARG after the removal step. As suggested in Mahmoudi et al. (2022), although the Kuhner move seems necessary and sufficient for good mixing when $\mu = 0$, it results in poor convergence for real data with mutations.

In our proposal (SGPR), we instead use threading to sample from the approximate posterior. This proposal now takes the data into account and is more likely to reach good likelihoods. In Section A.4.3, we show that when assuming that the threading algorithm approximately samples

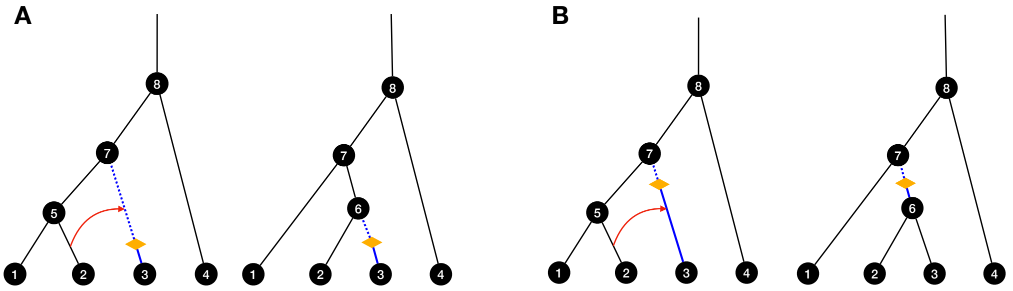

Figure S9: Similarities and differences between SGPR and ARGweaver proposals. When ARGweaver chooses branch $(3,7)$ to remove in the first tree, there are two possible branches to remove in the second tree, namely either $(3,6)$ or $(6,7)$. The two cases can be induced by having different locations of the cut on the same branch $(3,7)$ in the first tree (a lower cut in A versus a higher cut in B). Orange oval: a cut in the SGPR move; blue branches: affected branches; blue dashed parts: removed partial branches.

from posterior, the acceptance ratio will not contain the likelihood ratio term, which means we can introduce big moves in the proposal while having good acceptance probability.

We also point out that the removal scheme of SGPR and the Kuhner move can be seen as a clarification of the branch selection algorithm in ARGweaver (Figure S9). ARGweaver removes a sequence of branches by using a data structure called the "branch graph", and given the removed branch in the current tree, there can be up to two choices for removing a branch in the next tree. We illustrate with an example in Figure S9 that if we specify a location of the cut in the previous branch, then the next branch choice is always unique. If we do not know the location of the cut, then there can indeed be two possible next branches to choose from.

##### A.4.3 How to regraft the sub-graph to generate an updated ARG

After pruning a sub-graph, we need to regraft it to generate an updated ARG. In our proposal, we use our threading algorithm to re-thread the cut point onto the remaining graph $H$ after pruning the sub-graph, to obtain a new ARG state $G'$. This leads to a substantial improvement over the Kuhner move because the threading algorithm considers the data and will find good genealogies consistent with the observed allelic patterns. We show that with the following acceptance ratio, the MCMC will achieve detailed balance:

$$A_H(G \to G') = \min\left\{1, \frac{\mathbb{P}(G'|D)q_H(G' \to G)}{\mathbb{P}(G|D)q_H(G \to G')}\right\},$$

where $q_H(G \to G')$ denotes the probability of updating $G$ to $G'$ via the sub-graph pruning from $G$ to $H$ and re-grafting from $H$ to $G'$.

*Proof.* The target distribution of the MCMC is $\mathbb{P}(G|D)$ and we have

$$\mathbb{P}(G|D)\int_H q_H(G \to G')A_H(G \to G')dH$$

$$= \int_H \min\left\{\mathbb{P}(G|D)q_H(G \to G'), \mathbb{P}(G'|D)q_H(G' \to G)\right\}dH$$

$$= \mathbb{P}(G'|D)\int_H q_H(G' \to G)A_H(G' \to G)dH.$$

Hence, detailed balance is achieved with the stationary distribution being $\mathbb{P}(G|D)$. $\square$

Now, suppose the threading algorithm in the regraft step can approximately sample from the posterior distribution. Then,

$$\begin{aligned} q_H(G \to G') &= \mathbb{P}(H|G)\mathbb{P}(G'|D, H) \\ &= \mathbb{P}(H|G)\frac{\mathbb{P}(G'|D, H)}{\int_{G'':H\in\mathcal{H}(G'')}\mathbb{P}(G''|D, H)} \\ &= \mathbb{P}(H|G)\frac{\mathbb{P}(G', H|D)}{\int_{G'':H\in\mathcal{H}(G'')}\mathbb{P}(G'', H|D)} \\ &= \mathbb{P}(H|G)\frac{\mathbb{P}(G'|D)}{\int_{G'':H\in\mathcal{H}(G'')}\mathbb{P}(G''|D)}, \end{aligned}$$

where $\mathcal{H}(G'')$ denotes the set of all subgraphs of $G''$, and $G'' : H \in \mathcal{H}(G'')$ means the set of all ARGs that can reach $H$ after a sub-graph pruning, which is the same as the set of all ARGs that $H$ can reach after regrafting. The last equality follows since $H$ is a sub-graph of both $G'$ and $G''$. Using this proposal distribution, the acceptance ratio can then be calculated as

$$A_H(G \to G') = \min\left\{1, \frac{\mathbb{P}(H|G')}{\mathbb{P}(H|G)}\right\}.$$

Now, for the proposal scheme described in Section A.4.1, we have

$$\mathbb{P}(H|G) = \frac{1}{h(\Psi_x)},$$

where $h(\Psi_x)$ is the height of the tree $\Psi_x$ at the rightmost position $x$ of the previous cut.

The acceptance ratio now reduces to a very simple form, involving a ratio between the tree height before and after the proposal:

$$A_H(G \to G') = \min\left\{1, \frac{h(\Psi_x)}{h(\Psi'_x)}\right\}.$$

Consistent with intuition, we later show empirically that this ratio of tree heights is typically very close to 1 for sufficiently large sample sizes, meaning that few rejections will occur while making big updates to the ARG.

### B Simulation benchmarks

#### B.1 Comparison with PSMC

Since many ARG-based applications involve inferring pairwise coalescence times (Fan et al., 2022; Zhang et al., 2023; Wang and Coop, 2022), it is of interest to know whether using multi-sequence ARG inference methods leads to more accurate estimates than separately carrying out pairwise inference using PSMC models, especially extremely fast versions like XSMC (Ki and Terhorst, 2020) and Gamma-SMC (Schweiger and Durbin, 2023). Here, we investigate this question for Gamma-SMC with 50 sequences (the same setting as the constant population size scenario in Section 4.5 of the main text).

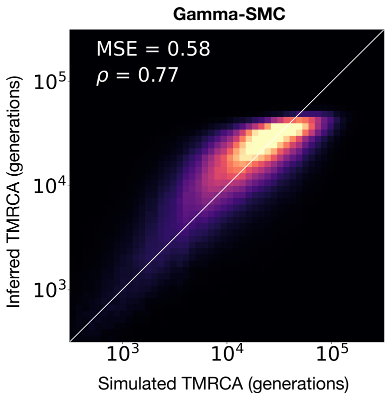

Figure S10: Performance of Gamma-SMC, an ultra-fast pairwise coalescent method, on inferring pairwise TMRCA for 50 sequences. The performance is similar to that of Relate, but worse than SINGER.

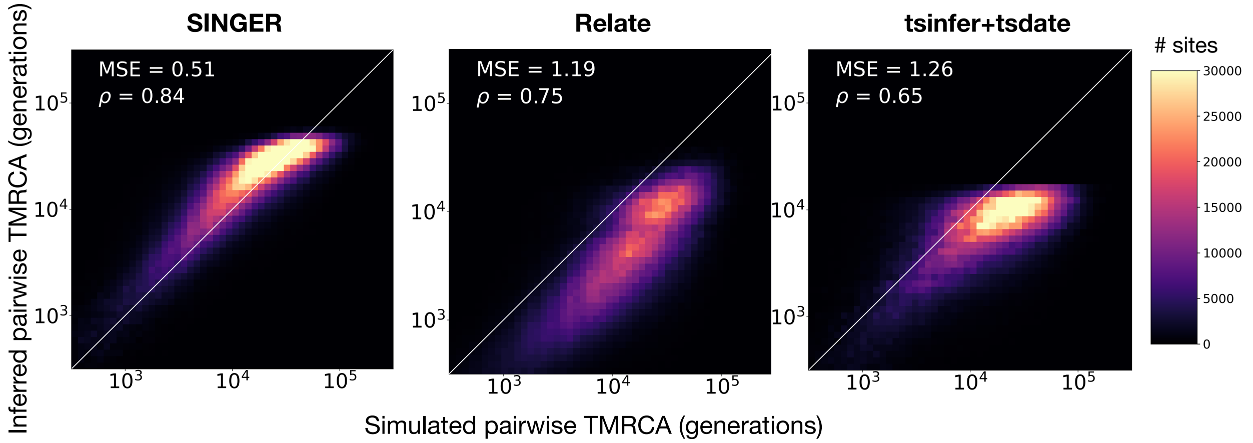

Figure S11: Robustness of ARG inference methods, when data is simulated with $N_e = 10,000$ but ARG is inferred with $N_e = 2,000$.

Surprisingly, despite modeling the joint genealogy of the entire sample, which should contain more information than just a pair of sequences, comparing Figure S10 with Figure 2A suggests that SINGER can substantially outperform Gamma-SMC, whereas Relate and tsinfer+tsdate do not seem to improve on Gamma-SMC in terms of MSE or correlation.

#### B.2 Robustness to model misspecification

To assess the robustness to model misspecification, especially that of population size history, we performed simulations with $N_e = 10,000$ but inferred ARGs assuming $N_e = 2,000$ (Figure S11). Further, we performed simulations under an inferred population size history for CEU (estimated using SMC++ (Terhorst et al., 2017) and available at https://github.com/PalamaraLab/ASMC_data/tree/main/demographies) and applied ARGs methods assuming a constant $N_e = 10,000$ (Figure 2B). For both scenarios, SINGER outperformed the other methods in terms of the accuracy of estimated pairwise coalescence times.

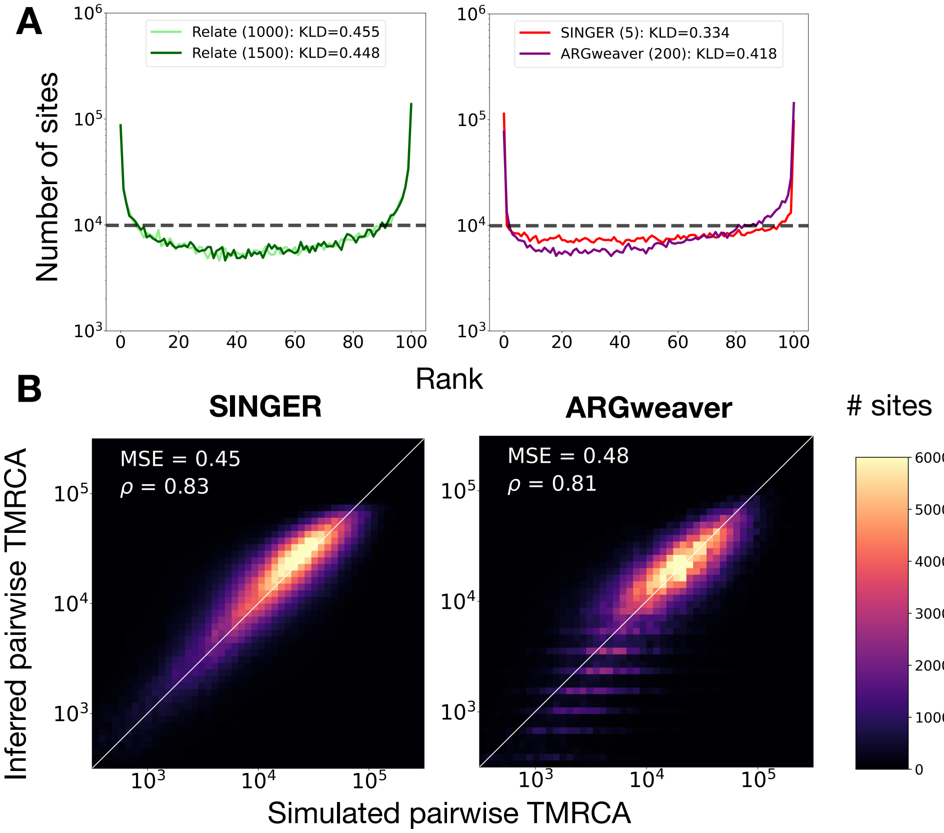

Figure S12: (A) Rank plots for different methods. In these results, considerably longer thinning intervals, indicated in parentheses, were used for ARGweaver and Relate. Relate is close to saturated thinning because changing the thinning interval from 1,000 to 1,500 results in little change in the rank plot. (B) Pairwise TMRCA inference from ARGweaver with thinning interval length 200 compared to that from SINGER with thinning interval length 5. ARGweaver still underperforms compared to SINGER in both rank plot and pairwise TMRCA, even with 40 times longer thinning than SINGER.

##### B.3 The impact of thinning on rank plots

Thinning interval can be an important parameter in MCMC samplers, as samples get less correlated with larger thinning intervals. In the rank plot analysis, one may argue that the U-shape is due to a lack of thinning. Here, we tried using 10 times larger thinning intervals (200 iterations) for ARGweaver than SINGER, and the suggested saturated thinning (1000 iterations) for Relate (Figure S12A).

All samplers deviated less from the uniform distribution with more thinning, but ARGweaver still performed worse in pairwise TMRCA, rank plot, and CI coverage compared to SINGER (Figure S12B), even with 10 times more iterations (which means hundreds to thousands times longer runtime!). Relate, however, seems to have reached saturated thinning, because the rank plots change little when thinning interval length is increased from 1000 to 1500. Still, the rank plots are not flat (Figure S12A). We believe this has to do with the fact that Relate does not sample tree topologies.

##### B.4 Fine-scale diversity estimation

In this section, we focus on the accuracy of estimating fine-scale diversity from ARG inference methods. By "fine-scale diversity", we mean average pairwise coalescence times (scaled by $4N_e\mu$) for small genomic windows. There are two ways of estimating this quantity: (1) by applying the classical nucleotide diversity estimator to the sequence data; or (2) by first inferring the underlying ARG and then calculating the average pairwise coalescence time in the ARG (scaled by $4N_e\mu$). We call them "SNP-based diversity" and "branch-length-based diversity", respectively.

In the main text, we use fine-scale diversity estimates to study local adaptation. Local adaptation refers to positive selection affecting a trait in a specific population. Due to hitch-hiking effects, we expect more recent coalescence times and reduced genetic diversity in the genomic regions that harbor recent positively selected genetic variants. Strong local reductions in genetic diversity are, therefore, often used to identify the genomic footprints of such selection. To accurately attribute the signal to a gene or regulatory element, genetic diversities need to be estimated at fine scales, e.g., 1 kb resolution. However, using SNP-based diversity can be very noisy at fine scales (Figure S13A), thereby limiting its utility. In general, for SNP-based diversity, there is a trade-off when choosing window sizes for diversity measures in which too small a window size causes high variance along the length of the genome, and too large window sizes causes over-smoothing where the signal of selection may be lost. However, if the ARG was directly observable, it would provide a much more precise measure of the reduction in diversity/coalescence times caused by positive selection. Effective estimation of local ARGs might, therefore, provide improved methods for identifying selective sweeps based on levels of genetic diversity. We illustrate this in Figure S13, where the ground truth from simulations is the branch-length-based diversity at each specific genomic coordinate. If posterior ARG samples from SINGER are used to obtain posterior averages of branch-length-based diversities, the agreement with the ground truth can be substantially improved (Figure S13D,E) compared to SNP-based estimates. In contrast, this benefit of using branch-length-based diversity estimates from inferred ARG is not as pronounced for Relate (Figure S13B), and even worse than SNP-based estimates for tsinfer+tsdate (Figure S13C).

#### C Applications to African data from the 1000 Genomes Project

##### C.1 Data and parameters for running SINGER

As mentioned in the main text, we analyzed 5 African groups (GWD,YRI, ESN, LWK and MSL) from the 1000 Genomes Project [4], with 40 genomes drawn from each group. SINGER requires an initial estimate of $N_e$ to run, for which we used the following simple estimator:

$$N_e \approx \frac{\pi}{4\mu},$$

where $\pi$ denotes the genome-wide average pairwise diversity, and $\mu$ the mutation rate per generation, per bp. Using $\mu = 1.2 \times 10^{-8}$, we obtained $N_e \approx 2 \times 10^4$. Because of the robustness of SINGER to model misspecification, this rough estimate was sufficient for us.

##### C.2 Large-scale diversity patterns

It has long been known that diversity levels are not constant along the genome, even on a rather large scale, e.g., 1 Mb (McVicker et al., 2009). Specifically, if we partition the genome into 1 Mb windows and compute their respective diversities, they will show up to 5-fold differences; this pattern is difficult to explain by a neutral model where demographic perturbations, in average, affects all

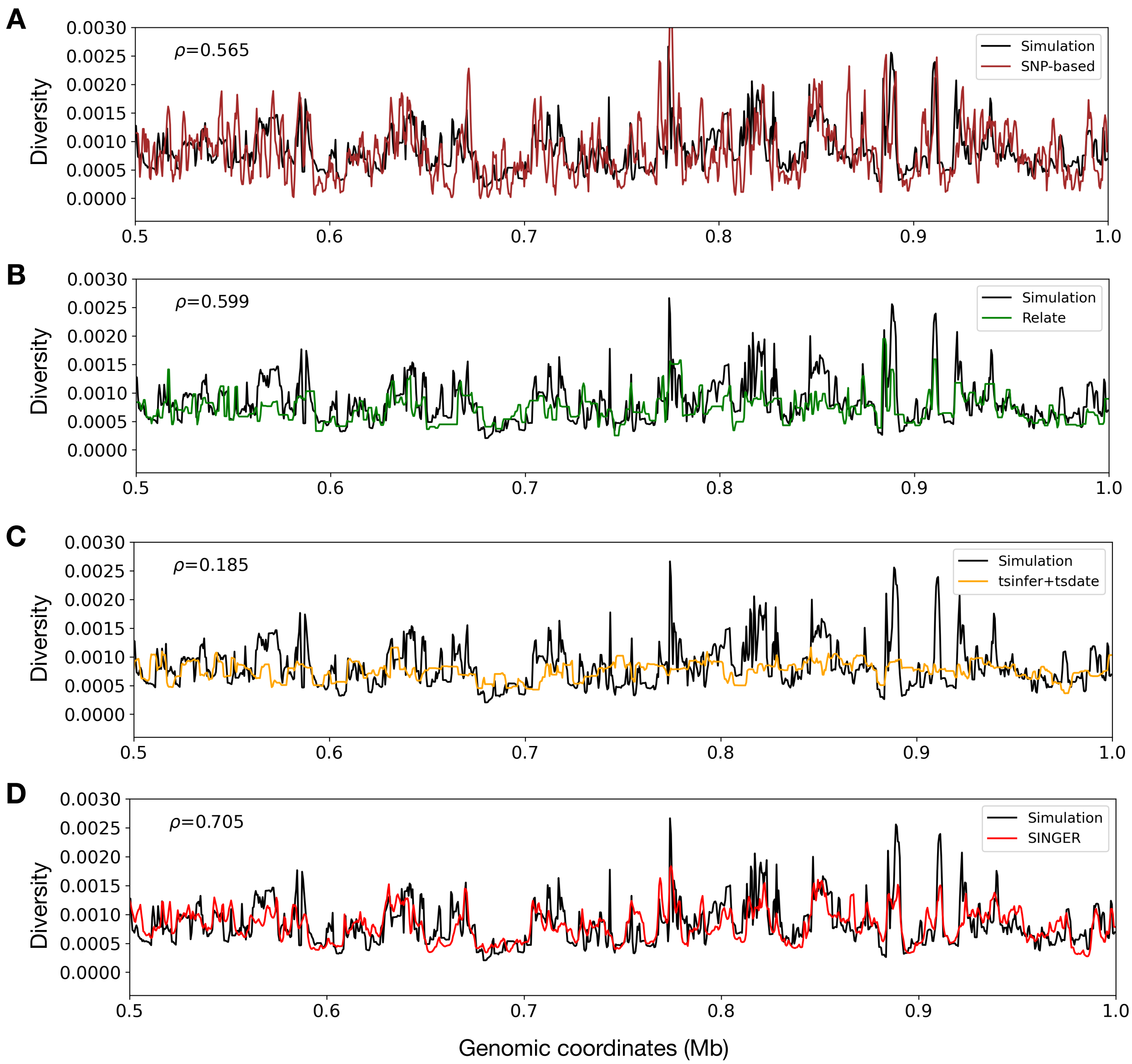

Figure S13: Different estimates of 0.5 kb fine-scale diversity against simulation ground truth from SNP-based sliding windows (A), Relate (B), tsinfer+tsdate (C), and SINGER (D).

genomic regions in the same way. A popular hypothesis proposed to explain this phenomenon is that background selection in the form of purifying selection acting on functional sites and affecting other sites through linkage, shapes the genome-wide diversity (McVicker et al., 2009; Murphy et al., 2022).

From the ARG inferred by Wohns et al. (2022) using tsinfer+tsdate, we extracted the sub-ARG for the same African individuals we considered in our study and compared it to the ARGs inferred by SINGER. In particular, for SINGER and tsinfer+tsdate, we calculated branch-length-based diversity estimates for each 1 Mb window, defined as the product of the mutation rate and the average pairwise distance in the ARG for the window, and compared them with the SNP-based diversity estimate from the VCF file. We observed that tsinfer+tsdate severely underestimated

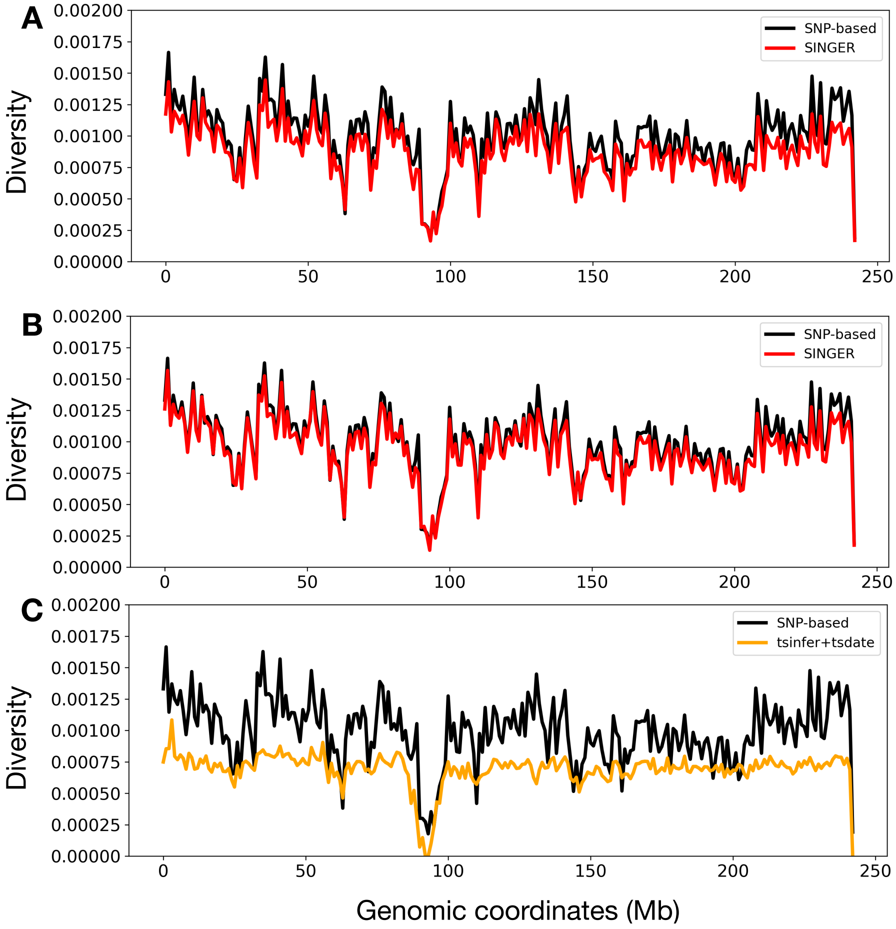

Figure S14: Diversity estimates in 1 Mb windows from SNP frequencies compared to (A) SINGER's initially inferred ARG, (B) SINGER's last ARG sample in MCMC, and (C) tsinfer+tsdate's inferred ARG.

genome-wide diversity levels and the extent of their variation, while SINGER yielded much closer fits (Figure S14). This suggests a better potential for analyzing background selection, and genome-wide patterns on variability in general, with SINGER.

For SINGER, it is interesting to note that the initial ARG sample does not fit the large-scale diversity pattern as well in some regions, but the fit gets substantially better with MCMC updates (Figure S14, Figure S15A). Besides accounting for the uncertainty in coalescence times and topology, this result demonstrates the benefit of performing MCMC rather than simply using the initial sample. We suggest inspecting such diversity plots as a part of the strategy for monitoring the convergence of the MCMC algorithm implemented in SINGER.

#### C.3 Convergence diagnostic of SINGER

To examine the convergence of the MCMC chain in SINGER, we looked at the fit of branch-length-based diversity predictions in inferred ARGs to SNP-based diversity in 1 Mb windows for real data. The deviation (measured as the mean squared error) decreased until a stable level was reached, indicating convergence (Figure S15A).

Additionally, we inspected the mutational mappings. Due to statistical phasing errors, gene

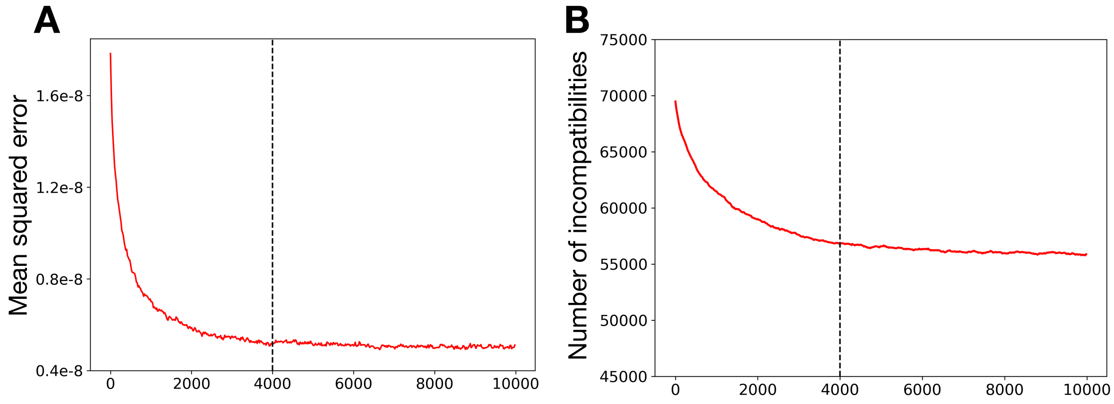

Figure S15: (A) A trace plot of deviation (measured in MSE) in diversity variations. (B) The number of incompatibilities for SINGER samples of Chr2. We used a burn-in of 4,000 iterations, when the chain appears to have reached stationarity.

conversions, sequencing errors, method imperfections etc, there will be some mutations that are not uniquely mappable to a single branch of a marginal tree, but the number should decrease with MCMC iterations (Figure S15B). We used a burn-in of 4,000 samples, after which the stabilization of both statistics suggested that the chain reached stationarity.

#### C.4 Coalescence distribution heatmaps used in introgression analysis

For the introgression analysis described in the main text, we inferred the site-specific distribution of pairwise coalescence times between a given leaf and all other leaves. However, the marginal distribution for a single tree is rather degenerate because it consists of a collection of point masses with different multiplicities (Figure S16A). In contrast, combining the trees sampled from the posterior using SINGER, which typically have different branch lengths and topologies, can substantially smooth this distribution (Figure S16B).

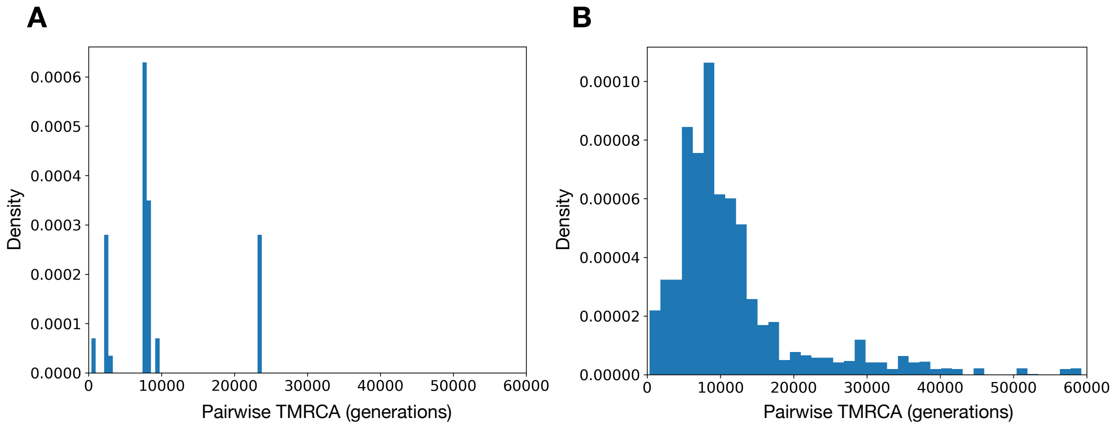

Figure S16: The coalescence distribution in a single marginal tree, when using a singer tree estimate (A) versus a collection of posterior samples (B) with different branch lengths and topologies. By sampling the branch length and topology uncertainties, we are able to obtain a much more stable distribution.

### D Additional Supplementary Figures

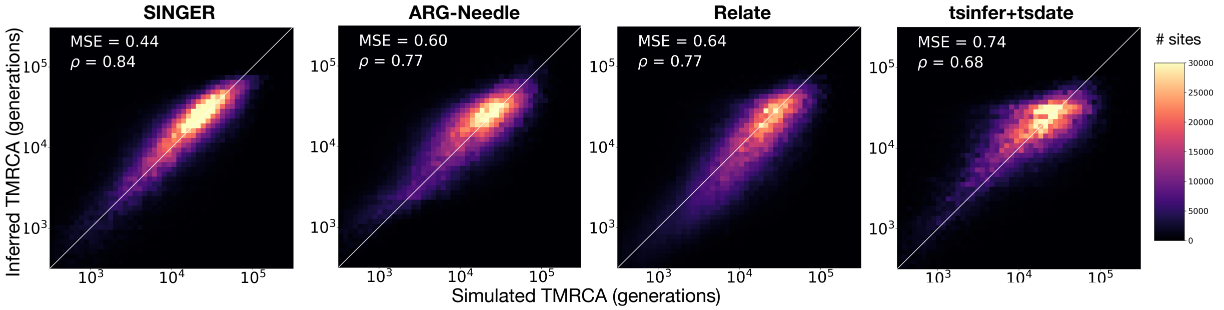

Figure S17: Performance of inferring pairwise TMRCA with SINGER, ARG-Needle, Relate and tsinfer+tsdate, with 300 sequences. SINGER performs the best, while ARG-Needle and Relate perform similarly, and tsinfer+tsdate performs the worst.

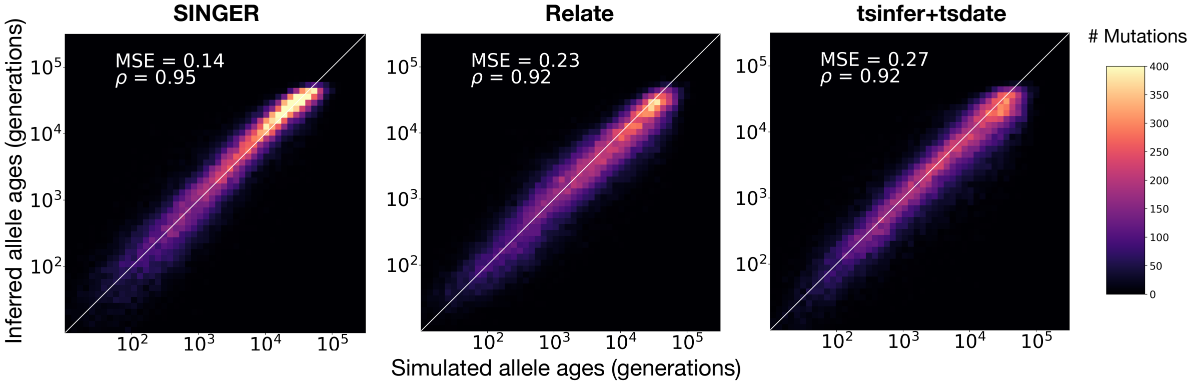

Figure S18: Performance of inferring allele ages with SINGER, Relate and tsinfer+tsdate, with 300 sequences. SINGER performs the best, followed by Relate, and tsinfer+tsdate performs the worst. ARG-Needle is excluded because it does not map mutations to branches of the inferred ARG.

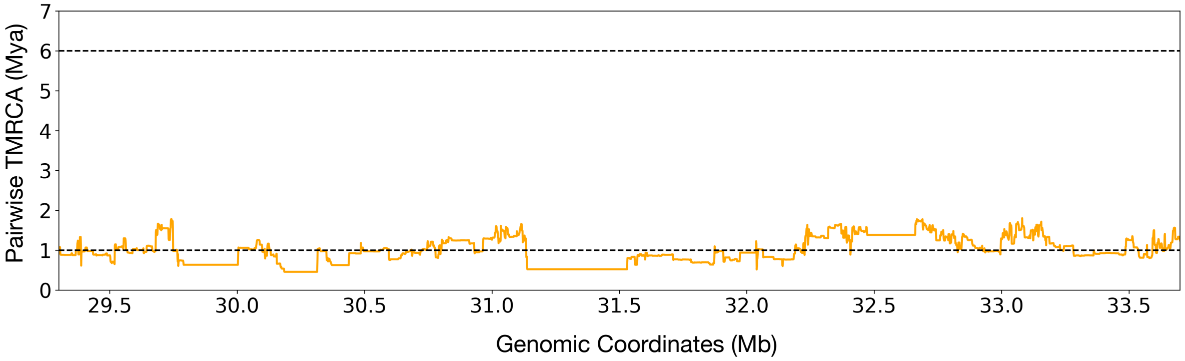

Figure S19: Average pairwise TMRCA in Africans inferred by tsinfer+tsdate from (Wohns et al., 2022). The 2 horizontal dashed lines are 1Mya and 6Mya, around African genome-wide average pairwise TMRCA and Human-Chimpanzee divergence time. Estimates from (Wohns et al., 2022) showed no evidence of trans-species polymorphism.
